# AAV delivered lysosome-targeting chimeras mediate sustained antibody depletion *in vivo*

**DOI:** 10.64898/2026.07.05.736665

**Authors:** Jonathan Lee Yang, Kang Yong Loh, Cindy R. Sandoval Espinoza, Dina Schuster, Sean A. Yamada-Hunter, Louai Labanieh, Elena Sotillo, Crystal L. Mackall, Karl Deisseroth, Carolyn R. Bertozzi

**Affiliations:** Department of Chemistry, Stanford University, Stanford, CA 94305; Sarafan ChEM-H, Stanford University, Stanford, CA 94305; Department of Bioengineering, Stanford University, Stanford, CA USA 94305; Department of Biochemistry, Stanford School of Medicine, Stanford, CA USA 94305; Center for Cancer Cell Therapy, Stanford Cancer Institute, Stanford University School of Medicine, Stanford, CA 94305; Parker Institute for Cancer Immunotherapy, Stanford University School of Medicine, Stanford, CA, United States of America; Weill West Coast Cancer Hub, Stanford, CA, United States of America; Department of Pediatrics, Stanford University School of Medicine, Stanford, CA, United States of America; Department of Medicine, Stanford University School of Medicine, Stanford, CA, United States of America; Stanford Cancer Institute, Stanford University School of Medicine, Stanford, CA, United States of America; Ludwig Center for Cancer Stem Cell Research and Medicine, Stanford University School of Medicine, Stanford, CA, United States of America; Department of Psychiatry and Behavioral Sciences, Stanford University, Stanford, CA, USA 94305; Howard Hughes Medical Institute, Stanford University, Stanford, CA 94305

## Abstract

Immunoglobulin G (IgG) is a critical effector of the adaptive immune system, but IgG autoantibodies against self-antigens drive pathology across a wide range of autoimmune diseases. Lowering circulating IgG is a validated and promising therapeutic avenue. Here we developed <u>g</u>enetically-<u>e</u>ncoded lysosome targeting <u>c</u>himeras (GELYTACs) that target circulating IgGs for clearance and degradation. The GELYTACs comprised two protein modules derived from insulin-like growth factor 2 (IGF2) and an IgG-binding nanobody, respectively, and mediated clearance of plasma IgG via the lysosomal trafficking receptor IGF2R. To achieve long-lasting IgG depletion, we encoded GELYTACs in an AAV gene therapy vector and established continuous expression in mice. We also developed conditional GELYACs that are activatable with disease-specific proteases or small molecule drugs and show that GELYTACs can be secreted from cellular therapies. This work establishes GELYTACs as a possible therapeutic modality that is deliverable using genetic medicine approaches.

**Summary References:**

1. Ahn, G., Banik, S. M. & Bertozzi, C. R. Degradation from the outside in: Targeting extracellular and membrane proteins for degradation through the endolysosomal pathway. Cell Chem. Biol. 28, 7, 1072–1080 (2021).
2. Ahn, G. et al., LYTACs that engage the asialoglycoprotein receptor for targeted protein degradation. Nat Chem Biol 17, 937–946 (2021).
3. Bunin, A. et al., Novel Bispecific Degrader BHV-1300 Achieves Rapid, Robust, Selective, and Transient IgG Reduction in Preclinical Models Including Nonhuman Primates, American Academy of Neurology (AAN) 2024 Annual Meeting.
4. Biohaven Reports Positive Degrader Data Achieving > 80% Sustained Reductions in Total IgG with Potential First-in-Class BHV-1300, Biohaven press release (2025).
5. Klein-Schneegans, A. S., Kuntz, L., Fonteneau, P. & Loor F. Serum Concentrations of IgM, IgGl, IgG2b, IgG3 and IgA in C57BL/6 Mice and their congenics at the lpr (lymphoproliferation) locus. J. Autoimmun. 2, 6, 869-875 (1989).
6. Monteys, A. M. et al. Regulated control of gene therapies by drug-induced splicing. Nature 596, 7871, 291–295 (2021).
7. Goertsen, D. et al. AAV capsid variants with brain-wide transgene expression and decreased liver targeting after intravenous delivery in mouse and marmoset. Nat Neurosci 25, 106–115 (2022).

## Main Text

Immunoglobulin G (IgG) is a central participant in adaptive immunity against foreign pathogens and cancers^1^. Dysregulated IgG expression is implicated in many autoimmune diseases such as systemic lupus erythematosus (autoantibodies against various nucleic acids)^2^, autoimmune encephalitis (autoantibodies against NMDA receptors in the brain)^3^, rheumatoid arthritis (antibodies against citrullinated proteins)^4,5^, and blood and organ transplant rejection^6,7^. In addition to autoimmune diseases, IgG overexpression and accumulation lead to cellular senescence, a key driver of aging in both mice and humans^8^.

The current standard of care for antibody-driven autoimmune diseases includes B-cell depletion^9^, plasma exchange^10^, inhibition of FcRn (receptors responsible for IgG recycling)^11^, steroids^12,13^, complement inhibitors^14^, cytokine blockers^15^ and JAK kinase inhibitors^16^. While these treatments improve the quality of life for patients with autoimmune diseases, they require lifelong regimens. Further, long term use of these therapies has serious side effects such as increased risk of infection^17,18^, cancer^19,20^ and blood clotting^21^. Therefore, new mechanisms to deplete IgG in a prolonged yet controlled manner warrant pursuit.

Extracellular targeted protein degradation (TPD) strategies are, in principle, well-suited to tackle the problem of IgG depletion^22,23^. Such technologies include <u>ly</u>sosome <u>ta</u>rgeting <u>c</u>himeras (LYTACs)^24,25^ and related platforms^26,27^ that comprise target binding moieties chemically conjugated to ligands that bind lysosomal trafficking receptors. These bifunctional molecules have been designed to engage widely expressed receptors such as the insulin-like growth factor 2 receptor (IGF2R, also known as the mannose-6-phosphate receptor), or tissue-specific receptors like the asialoglycoprotein receptor (ASGPR). Prior work has shown that extracellular degraders administered to animals^28^ or humans^29^ can effectively deplete circulating immunoglobulins while the drug is on board. In a phase 1 clinical study, an IgG-depleting LYTAC-like peptide conjugate maintained reduced levels of circulating IgG during a course of weekly drug injections^30^. These precedents point to the LYTAC mechanism as a promising approach to reduce circulating IgG levels for autoimmune disease patients. However, they require frequent and indefinite drug administration. We considered an alternative strategy in which therapeutic LYTAC molecules are continuously produced *in vivo* via gene therapy. Toward this end, we recently reported the development of genetically-encoded LYTACs (GELYTACs) comprising a target-binding nanobody and an engineered IGF2 protein that binds IGF2R^31^. Here, we applied this principle to the development of IgG-depleting GELYTACs, including variants subject to temporal control with enzymatic or small molecule switches. We further show, using a model target, that GELYTACs can be secreted from CAR-T cells without compromising their cytotoxicity. We achieved continuous production of GELYTACs in mice using an AAV8 vector that drives expression across a broad range of tissues. Mice treated with one dose of this gene therapy showed prolonged (2 month) reduction in steady-state IgG1, IgG2b, and IgG2c levels of >85% when compared to mice treated with AAV8 encoding for IgG-targeting nanobody only. We then show that AAV GELYTACs could clear rheumatoid arthritis causing antibodies, which potentially results in improved disease outcomes when compared to treatment with AAV8 encoding for IgG-targeting nanobody only.

## Results

### Design of IgG-depleting GELYTACs that engage murine IGF2R

Our earlier GELYTACs were designed to engage human IGF2R with an engineered human IGF2 module. To test GELYTACs in a gene therapy platform using murine models, we switched to the <u>m</u>urine isoform of IGF2 (mGELYTACs). We generated four mGELYTAC constructs varying in orientation and linker length and narrowed our design to include, from N-to C-terminus: an epitope tag, murine IGF2, a 20-amino acid glycine-serine linker, and a nanobody against IgG (Fig. 1a and Extended Data Fig. 1a, b). Unless otherwise noted, mGELYTAC constructs were situated under the control of an EF1-α promoter.

**Fig. 1:**
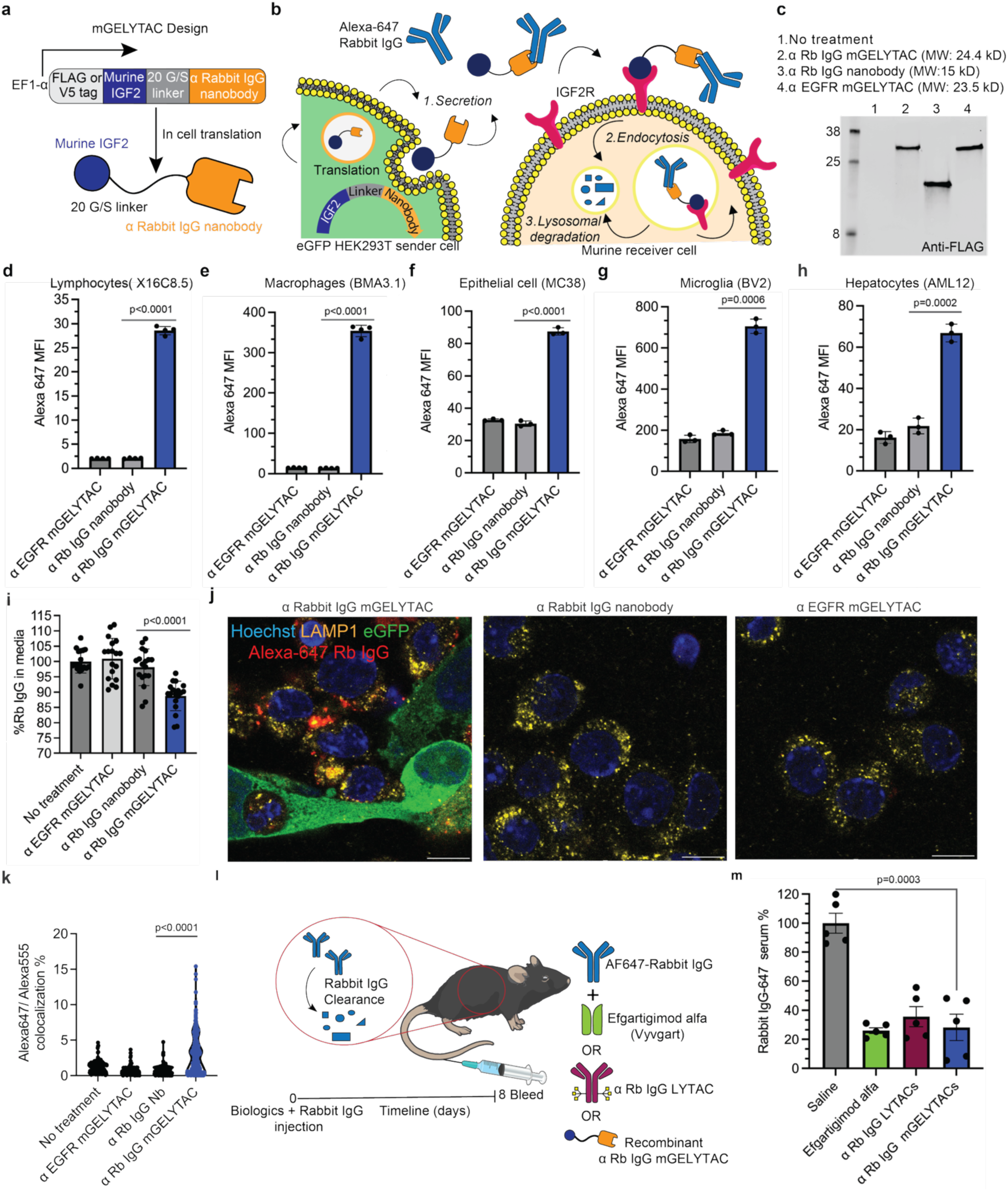
mGELYTAC mediate uptake IgG in murine cell culture & *in vivo*. **a**, Schematic of murine genetically-encoded LYTACs (mGELYTACs), bifunctional proteins consisting of a binder (i.e. nanobody) to IgG and murine IGF2, a 7.5 kDa protein that binds to IGF2R. **b**, Schematic of co-culture experiments depicting mGELYTAC secretion from sender cells mediating internalization of Alexa-647 labeled IgG into receiver cells. **c**, α FLAG western blot of conditioned supernatant from HEK293T cells transiently transfected with α Rabbit IgG mGELYTACs, α Rabbit IgG nanobody only, and a non-targeting mGELYTAC control (targeting human EGFR). **d-h**, Alexa-647 MFI (median fluorescence intensity) values from a variety of murine cell lines representing different cell types cocultured with eGFP HEK293T cells secreting α rabbit IgG mGELYTAC, rabbit-IgG nanobody control, or a control mGELYTAC (targeting human EGFR). Cocultured cells were treated with Alexa-647 rabbit-IgG (100 nM) for 24 hours. Then, the cell mixture was analyzed using flow cytometry. Error bars represent the SD from 3-4 biological replicates. p values were determined using Welch’s t-test. **d**, X16C8.5 lymphocytes. **e**, BMA3.1 macrophages. **f**, MC38 epithelial cells. **g**, BV2 microglia cells. **h**, AML12 hepatocyte cells. **i**, To quantify Alexa-647 rabbit-IgG (25 nM) clearance in the coculture system, media was analyzed for Alexa-647 fluorescence 72 hours after Alexa-647 rabbit-IgG addition using a plate reader. This experiment was repeated 18 times. Error bars represent the SD from 18 biological replicates. p values were determined using Welch’s t-test. **j**, Confocal fluorescence images of cocultured eGFP HEK23T cells secreting α rabbit IgG mGELYTAC, rabbit-IgG nanobody control, or a control mGELYTAC (targeting human EGFR) co-cultured with murine J77A macrophages. Cocultured cells were treated with Alexa-647 rabbit-IgG (100 nM) for 24 hours and stained with LAMP1 lysosome marker and Hoechst (nuclear stain). Scale bars, 10 μm. **k**, Quantification of colocalization of Alexa-647 rabbit-IgG with Lamp1 staining in confocal images (**j** and Extended Data Fig. 1e). The quantification is defined as pixels within the image that contain both the Alexa-647 (rabbit IgG) and Alexa-555 (LAMP1) signal. Error bars represent the SD from at least 114 cells. p value determined using Mann-Whitney U non-parametric test. **l**, Schematic outlining comparison experiment between recombinant mGELYTAC and other IgG depletion therapeutics that clear Alexa-647 tagged rabbit IgG *in vivo*. **m**, Plasma Alexa-647 rabbit IgG levels at day 8 for mice treated with 15 mg/kg Alexa-647 rabbit IgG in addition to saline, efgartigimod alfa (10 mg/kg), α Rb IgG LYTACs (15 mg/kg), or α Rb IgG mGELYTAC (3.1 mg/kg).

For initial proof-of-concept, we designed mGELYTACs to bind rabbit IgG using a rabbit IgG-specific nanobody^32^. We tested mGELYTACs in cell culture using a sender-receiver model, wherein mGELYTACs were transiently expressed in and secreted from eGFP-HEK293T “sender” cells (Fig. 1b). Secreted mGELYTACs can bind to an Alexa-647-labeled rabbit IgG in the media and direct its internalization into murine “receiver” cells. Once trafficked to the lysosome of the receiver cell, the rabbit IgG protein would be degraded, but not the Alexa-647 dye^33^. Thus, we can use fluorescent readouts to monitor mGELYTAC-mediated IgG uptake by the fold-change of Alexa-647 fluorescence.

We confirmed secretion of mGELYTAC or control proteins (control mGELYTAC^34^ and rabbit IgG-specific nanobody) into the media of transfected eGFP-HEK293T cells by α FLAG western blotting (Fig. 1c). Using flow cytometry, we showed that mGELYTACs mediate the uptake of Alexa-647-labeled rabbit IgG by murine cell lines derived from various tissues, including lymphocytes (X16C), macrophages (BMA3.1, J77A, RAW264.7), epithelial cells (MC38), microglia (BV2), and hepatocytes (AML12). Receiver cells co-cultured with mGELYTAC-secreting sender cells exhibited significant uptake of Alexa-647-labeled rabbit IgG when compared to controls (Fig. 1d-h and Extended Data Fig. 1c,d). We also investigated mGELYTAC’s ability to clear its target from the media of cell co-cultures. We incubated murine X16C.8 lymphocytes alongside eGFP-HEK293T cells secreting mGELYTACs specific for rabbit IgG. Alexa-647 rabbit IgG added to the supernatant was found to be depleted after 72 hours (Fig. 1i).

To confirm lysosomal trafficking of the mGELYTAC-Alexa-647-rabbit IgG complex, we performed imaging studies with J77A murine macrophage cells cocultured with mGELYTAC– (or control protein)-secreting eGFP-HEK293T cells in media containing Alexa-647-rabbit IgG. We observed Alexa-647 colocalization with the lysosome marker LAMP1 (Fig. 1j and Extended Data Fig. 1e), indicating rabbit IgG was internalized into the lysosomes (Fig. 1k). Staining for mGELYTAC directly with α FLAG antibodies confirmed its colocalization with rabbit IgG (Extended Data Fig. 1f,g). Lastly, to confirm that IgG uptake was dependent on IGF2/IGF2R binding, we performed a receptor competition experiment. Addition of recombinant murine IGF2 to the co-culture media caused near complete suppression of Alexa-647 uptake by the receiver cells, suggesting that mGELYTAC-IGF2R binding is required for rabbit IgG internalization (Extended Data Fig. 1h).

Lastly, to test if mGELYTAC protein is functional *in vivo*, we injected Alexa-647 labeled rabbit IgG with recombinant α rabbit IgG mGELYTAC or other validated IgG depleting therapeutics (α rabbit IgG LYTACs^25^ or FDA approved Fc inhibitors (Efgartigimod alfa^35^)) into mice and monitored plasma Alexa-647 signal at the 8-day mark (Fig. 1l, Extended Data Fig. 1 i-l). Compared to saline treated mice, mGELYTACs mediated significant depletion of AF647-rabbit IgG, and achieved similar levels of IgG clearance compared other validated IgG degraders (Fig. 1m).

### mGELTACs deplete exogenously added IgGs in vivo

Having shown that mGELYTACs mediate the uptake and clearance of Alexa-647-rabbit IgG in cell culture, and when directly injected in vivo, we next tested their ability to do so when produced and secreted *in vivo*. To accomplish this, we generated an AAV8 construct encoding the rabbit IgG-specific mGELYTAC or control genes. When injected into the murine tail vein, AAV8 typically infects many cell types (most prominently liver hepatocytes^36^) and drives long-term transcription and translation of the transgene^37,38,39^ (Fig. 2a). We injected AAV8 encoding for α rabbit IgG mGELYTAC or control proteins into the tail vein of C57BL/6 mice, and isolated plasma via submandibular bleed every 3 days until 15 days post injection (Fig. 2b). Western blot analysis of these samples showed a gradual increase in plasma protein levels at least until day 15 (Fig. 2c).

**Fig. 2:**
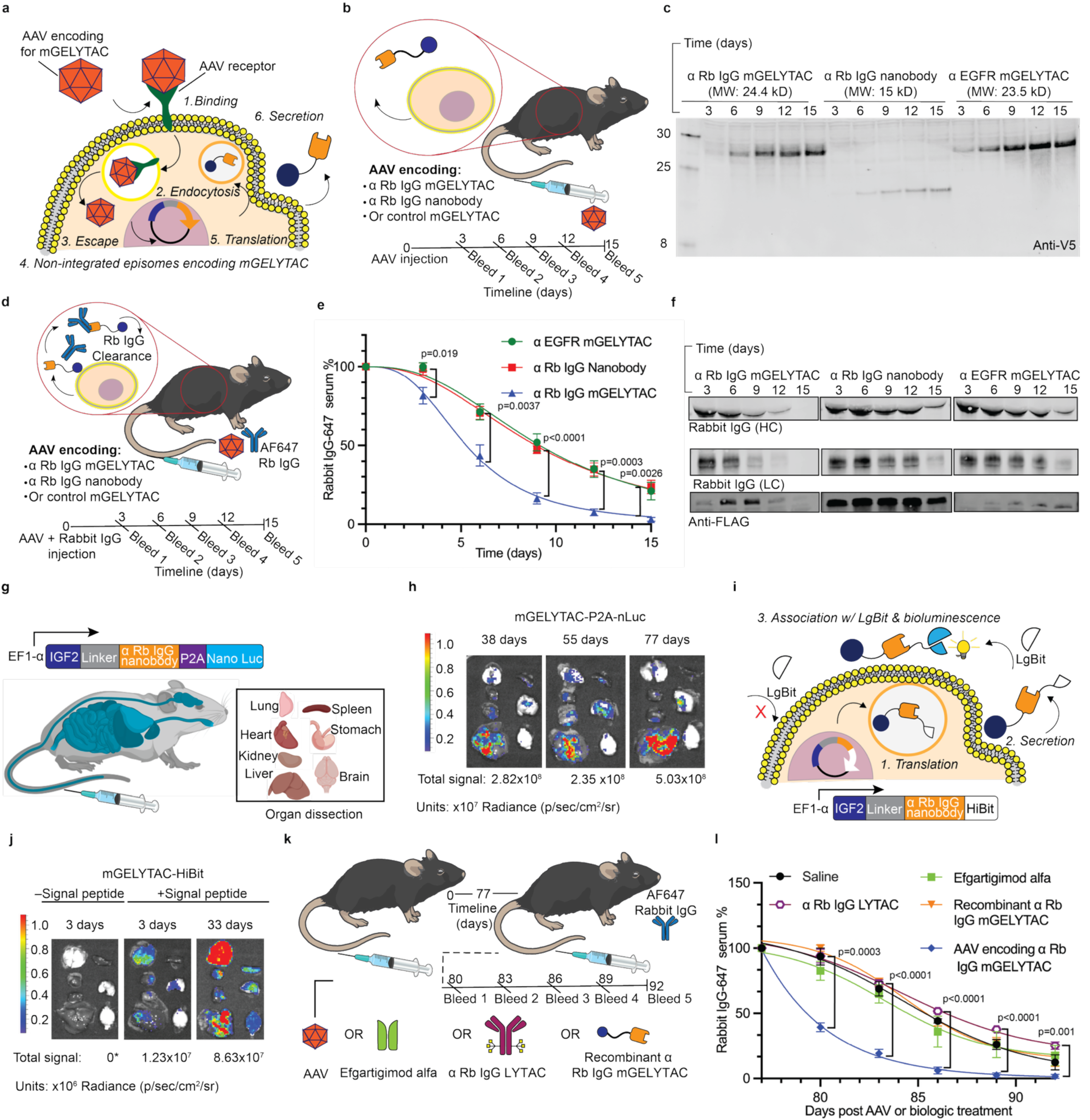
Gene therapy delivered mGELYTACs clear exogenous IgGs *in vivo.* **a**, Schematic of AAV8 encoding for mGELYTAC infecting a host cell. **b**, Schematic outlining AAV infection experiment. AAV viral particles encoding for rabbit IgG targeting mGELYTACs were injected via tail vein. Submandibular bleeds were performed every 3 days from day 3-15 and then analyzed via α V5 western blot. **c**, α V5 western blot of murine plasma collected from C57BL/6 mice infected with AAV8 encoding for rabbit IgG mGELYTAC, rabbit IgG nanobody, or control mGELYTAC human EGFR mGELYTAC^34^. **d**, Schematic outlining exogenous IgG clearance in vivo. AAV8 particles encoding for mGELYTACs or controls were mixed with 37.5 mg/kg of Alexa-647 labeled IgG and injected via tail vein. Plasma was collected every 3 days until day 15 where the Alexa-647 IgG levels were analyzed either via plate reader (**e**) or western blot (**f**). **e**, Alexa-647 rabbit IgG serum clearance as a function of time for mice treated with AAVs encoding for rabbit IgG (triangles), rabbit IgG nanobody (squares), or control mGELYTAC (targeting human EGFR, circles). Rabbit IgG clearance was measured via Alexa-647 plate reader assay on samples collected between 3-15 days. Error bars represent the SEM from 9 biological replicates for rabbit IgG mGELYTACs, 8 biological replicates for rabbit IgG nanobody, and 7 biological replicates for control (human EGFR) mGELYTACs. p values were determined using Welch’s t-test. **f**, Western blot analysis of Alexa-647 rabbit IgG levels determined via IR700 measurement. Labeled heavy chains appear at ∼50 kDa and labeled light chains appear at ∼25 kDa. α FLAG staining was also performed to analyze mGELYTAC or control levels corresponding to exogenous rabbit IgG levels. **g**, Schematic for mGELYTAC containing nano luciferase downstream to enable bioluminescence imaging and arrangement of organs post dissection. The organ arrangement schematic represents the order organs in all organ imaging in this figure. **h**, A comparison of bioluminescence from organs from mice treated with AAV8 encoding mGELYTAC-P2A-nLuc at 38 days, 55 days, or 77 days. **i**, Schematic depicting extracellular mGELYTAC detection. mGELYTACs fused to HiBit (split nano Luc fragment) are secreted from cells, where it reconstitutes with LgBit to form an intact and functional nano luciferase. mGELYTAC-HiBit in the secretory pathway does not give bioluminescence because LgBit is too large to pass through membranes. **j**, A comparison of bioluminescence from organs of mice treated with AAV8 encoding mGELYTAC-HiBit at days 3 and 33 post-injection and subsequently treated with LgBit and furimazine. The negative control is with AAV8 encoding mGELYTAC-HiBit without a signal peptide. *Indicates that the quantified bioluminescence was below 0; 0 is written because it is the closest logical value. **k**, Schematic outlining experiment to monitor AAV delivered mGELYTAC longevity in comparison with other validated IgG degrading therapeutic modalities (Efgartigimod alfa or α Rb IgG LYTAC) or recombinant α Rb mGELYTAC. C57BL/6 mice were injected with AAV viral particles encoding for rabbit IgG targeting mGELYTACs or other biologics. 77 days later, mice were injected with 15 mg/kg Alexa-647 labeled rabbit IgG, and submandibular bleed was performed every 3 days from day 80-92 and then analyzed via plate reader or western blot. **l**, AF647-rabbit IgG serum clearance as a function of time for mice treated with AAV encoding for α rabbit IgG mGELYTACs (rhombuses), recombinant α rabbit IgG mGELYTAC (triangles), Efgartigimod alfa (squares), α rabbit IgG LYTAC (hexagons), or saline (circles). Rabbit IgG clearance is measured via Alexa-647 plate reader assay on samples collected between 72-84 days. Error bars represent the SEM from 3-4 biological replicates. p values were determined using Welch’s t-test.

To assess rabbit IgG clearance activity, we co-injected AAV8 viral particles encoding rabbit IgG-targeting mGELYTAC (or controls) along with 0.75 mg (37.5 mg/kg) of rabbit IgG. C57BL/6 mice have ∼1 mg of total endogenous IgG2b, the most abundant murine IgG isotype, making our rabbit IgG dose physiologically comparable^40^. We then took bleeds every 3 days until day 15 to assess target clearance by detecting Alexa-647 fluorescence in a plate reader assay (Fig. 2d). We observed a ∼2-fold increase in the rate of clearance of rabbit IgG in the presence of mGELYTAC compared to non-targeted control mGELYTAC (Fig. 2e). A Western blot detecting Alexa-647-rabbit IgG further validated the clearance trend observed by the plate reader assay (Fig. 2f). Additionally, by blotting for mGELYTAC in the plasma, we observed a correlation between levels of mGELYTAC and its target rabbit IgG. Notably, mGELYTAC was detected at higher levels when its target was present but tapered off as rabbit IgG was cleared (Fig. 2f). This suggests that binding to target IgGs in serum stabilizes mGELYTACs, and that once the target clears, mGELYTACs are more rapidly cleared out of circulation. Lastly, we observed that IgGs are largely concentrated in liver and lungs post AAV8 treatment (Extended Data Fig. 2a).

Following verification of AAV8-delivered mGELYTACs efficacy *in vivo*, we studied the origin and durability of mGELYTAC production. We engineered a construct encoding the mGELYTAC fused to nano-luciferase with an intervening self-cleaving peptide (P2A) (Fig. 2g). To look at mGELYTAC expression over time, mice were treated with the nano-luciferase substrate furimazine at 38, 55, and 77 days. Mouse organs were harvested and imaged using an IVIS system (Fig. 2h). Bioluminescence signal generated by luciferase acting on furimazine served as a proxy for mGELYTAC expression. We observed stable expression from all collected organs (aside from the brain) with the liver being the main producer. To show that brain specific transduction is possible, we showed that retroorbital delivery of a brain penetrating AAV serotype (AAV.PhP.eb) incorporating a neuron-specific promoter (hSyn) resulted in brain expression of mGELYTAC (Extended Data Fig. 2b,c).

Next, we identified the tissues the mGELYTACs localize to after secretion. To distinguish secreted mGELYTACs in tissues from intracellular mGELYTACs in the secretory pathway, we utilized a split nano-luciferase assay (Fig. 2i). We fused an 11 amino acid fragment of nano-luciferase (HiBit) to the C-terminus of mGELYTAC and injected the complementary non-cell permeable fragment of nano-luciferase (LgBit) along with furimazine during analysis. Following verification in cell culture (Extended Data Fig. 2d), we tested this system *in vivo* and observed accumulation of mGELYTAC in all tissues with the lungs having the highest signal (Fig. 2j). To verify that the signal is coming from extracellular mGELYTAC, we also included a control construct lacking the signal peptide and observed no detectable bioluminescence (Fig. 2j).

Successively, we examined if the rise in detectable mGELYTACs corresponding to the introduction of exogenous IgG in serum (Fig. 2f) is consistent in organs. We utilized the same HiBit/LgBit assay, but this time also injected rabbit IgG. At the 6-day mark, we observed elevated bioluminescence in the lungs and hearts of rabbit IgG treated mice compared to organs from mice not given the rabbit IgG (Extended Data Fig. 2e).

The main advantage of AAV-delivered mGELYTACs over traditional IgG depleting biologics is its long window of activity (Extended Data Fig. 2f-j). To show this, we first injected mice with saline, AAV8 encoding for mGELYTAC, efgartigimod alfa, LYTAC, or recombinant mGELYTACs. At 77 days post treatment, mice were then injected with Alexa-647 labeled rabbit IgG followed by fluorescent analysis of plasma collected at days 80-92 (Fig. 2k). Out of all the treatments, only the AAV8 delivered mGELYTAC displayed significant levels of rabbit IgG depletion compared to the saline control despite having been introduced into the mice over 2 months prior (Fig. 2l, Extended Data Fig. 2k). Furthermore, prior mGELYTAC-AAV8 infection followed by target IgG injection resulted in more efficient target clearance compared to simultaneous gene therapy treatment and target administration. Presumably, in this paradigm the mGELYTAC protein has time to accumulate in serum to more efficacious levels.

Introducing non-native proteins like GELYTACs into animals brings the risk of α drug antibody (ADA) formation^41^. To test whether mice generate antibodies against AAV-delivered mGELYTACs, we treated mice as above then collected plasma after 55 days, a sufficient amount of time for a robust immune response in mice (Extended Data Fig. 2l)^42^. We tested the plasma for α mGELYTAC antibodies using a sandwich <u>e</u>nzyme linked immuno<u>s</u>orbent <u>a</u>ssay (ELISA) (Extended Data Fig. 2m) and included a murine α FLAG positive control since mGELYTACs contain a FLAG tag. Promisingly, no detectable α mGELYTAC antibodies were detected in plasma from infected mice (Extended Data Fig. 2n), suggesting a low immunogenic risk arising from circulating GELYTACs.

In addition, IGF2 fusion proteins may influence endogenous IGF2 clearance pathways. To test this, we measured the clearance of Alexa-647 labeled IGF2-MSA (mouse serum albumin) in mice treated with and without AAV8 encoding mGELYTAC and observed no significant differences in IGF2-MSA clearance (Extended Data Fig. 2o-r).

### Synthetic biology applications of GELYTACs

Next-generation gene therapy technologies are integrating mechanisms to control expression or activity of the transgene in a tissue-specific or temporally-controllable manner^43,44^. For example, protease-activatable antibodies^45^ or CAR-T cells^46^ have been designed for selective activation in tumor microenvironments. Small molecule switches have been designed for drug-dependent gene expression^44^. Finally, cell type-specific promoters have been integrated into gene therapy platforms to spatially restrict gene expression. We explored deployment of these three control mechanisms in the context of GELYTACs.

#### Protease-activatable GELYTACs

To design a protease-triggered mGELYTAC, we introduced an IGF2 masking protein into the mGELYTAC construct, separated from the active mGELYTAC with an intervening protease cleavage site. In this design, the mGELYTAC is caged by the IGF2 masking element and uncaged upon proteolytic cleavage (Fig. 3a). To identify suitable IGF2 masking moieties, we looked to the six known soluble IGF1/2 binding proteins (IGFBPs) (Fig. 3b)^47^. IGFBPs bind IGF2 with high affinities (low nM K_d_ values) and act endogenously to increase IGF2’s metabolic stability^48,49^. We expressed in HEK293T cells secreted mGELYTACs fused to various IGFBPs (IGFBP1-6) linked by a TEV protease cleavage site. All constructs exhibited full cleavage after treatment with 500 nM TEV protease (Fig. 3c and Extended Data Fig. 3a). To test the efficacy of TEV protease-gated mGELYTACs, we cocultured eGFP HEK293T cells secreting the aforementioned gated mGELYTACs with MC38 receiver cells. The coculture was supplemented with 185 nM Alexa-647-labeled rabbit IgG, and after 24 hrs, 500 nM TEV protease (or vehicle) was added. Flow cytometry analysis 24 hrs later revealed substantial uptake of Alexa-647-labeled rabbit IgG mediated by TEV protease-cleaved, but not uncleaved, mGELYTACs (Fig. 3d). In the control where IGFBP was replaced by mCherry, there is still a difference in rabbit IgG uptake between the with or without TEV protease which is likely attributed to mCherry providing steric hindrance. However, there is higher significant background without protease (∼3-fold increase), indicating that IGFBP binding to IGF2 further reduces background activity.

**Fig. 3:**
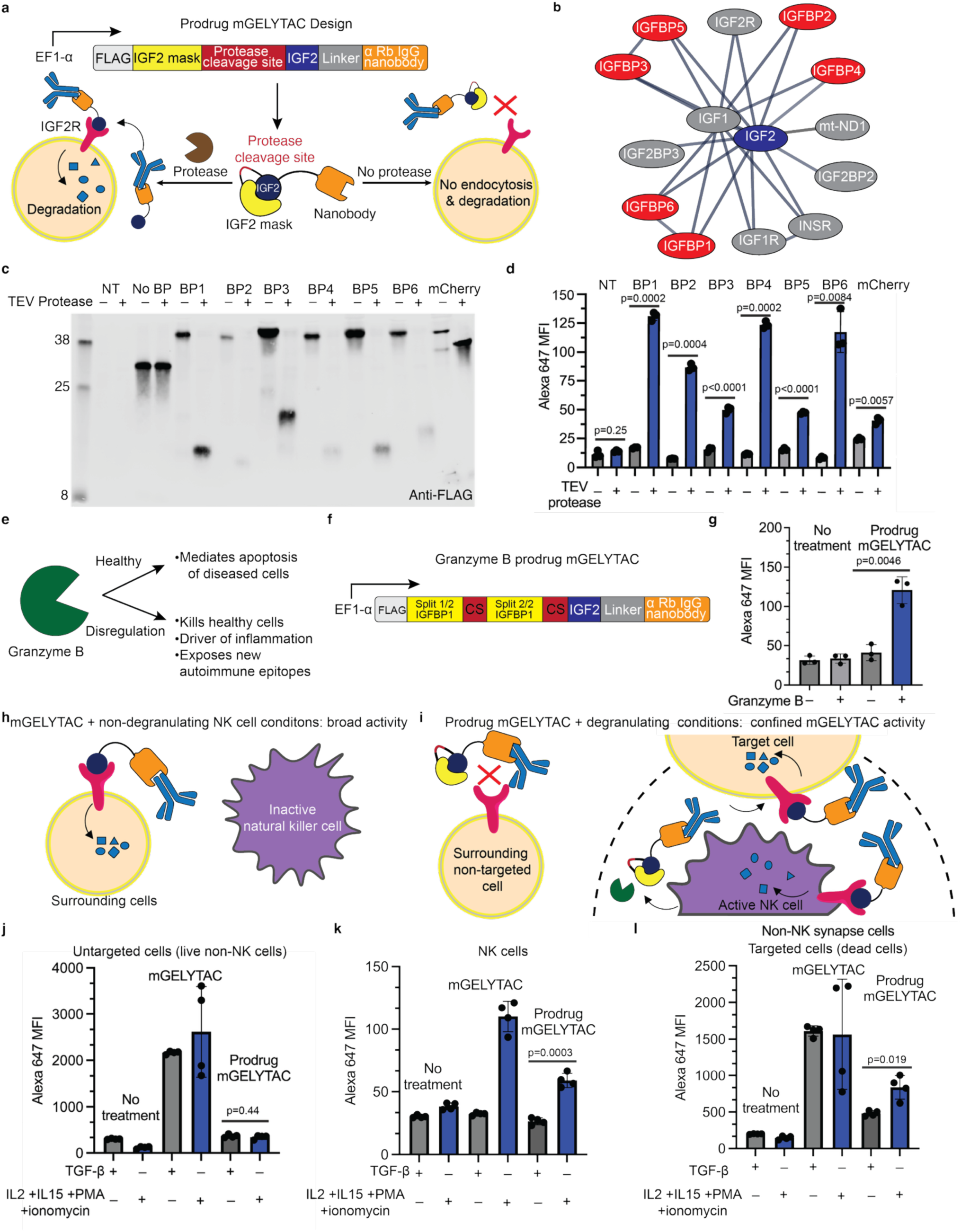
Development of protease-activatable mGELYTACs. **a**, Schematic outlining prodrug mGELYTAC design. mGELYTACs targeting rabbit IgG are genetically fused to an IGF2 mask with a protease cleavage site within the linker between IGF2 and the IGF2 mask. **b**, Murine IGF2 interactome^40^. **c**, α FLAG western blot of conditioned supernatant from HEK293T cells transiently transfected with a series of α Rabbit IgG prodrug mGELYTACs bearing IGF-binding proteins 1-6, control prodrug mGELYTAC bearing mCherry (Extended Data Fig. 3a), or no treatment (NT). After transfection, cells were either left untreated or treated with 500 nM TEV-protease. **d**, Alexa-647 MFI (median fluorescence intensity) values from MC38 cells cocultured with HEK293T cells secreting α rabbit IgG prodrug mGELYTACs with IGFBPs ranging from 1-6, control prodrug mGELYTAC (bearing mCherry instead of IGFBP), or no treatment (NT). After an initial 24 hr Alexa-647 rabbit IgG incubation, the cell mixture was either treated with 500 nM TEV-protease or left untreated for an additional 30 hrs before flow cytometry analysis. Statistics determined using Welch’s t-test. **e**, Schematic showing the role of granzyme B (GzmB) in healthy and diseased states. **f**, Schematic outlining the design of prodrug mGELYTAC gated by GzmB. **g**, Alexa-647 MFI (median fluorescence intensity) values from MC38 cells treated with supernatant from HEK293T cells transfected with or without GzmB gated prodrug mGELYTAC. In the positive GzmB condition, conditioned media was supplemented with 375 nM recombinant GzmB. p value determined using Welch’s t-test. **h**, Schematic outlining effects of non-gated mGELYTAC, which mediates universal IgG uptake. **i**, Schematic showing hypothesized GzmB prodrug mGELYTAC activity, where IgG uptake occurs in areas where there is high effective GzmB concentration (i.e. immune synapse). **j**, Alexa-647 MFI (median fluorescence intensity) values from live eGFP HEK293T transfected with or without GzmB gated prodrug mGELYTAC cocultured with NK92 cells (24 hrs). NK92 cells were either treated with TGF-β (immunosuppressive) or IL2 + IL15 + ionomycin + PMA (immune activating). Error bars represent the SD from 4 biological replicates. p values were determined using Welch’s t-test. **k**, Alexa-647 MFI values from live NK92 cells cocultured with eGFP HEK293T cells transfected with or without or GzmB gated prodrug mGELYTAC (24 hrs). NK92 cells under immunosuppressive and activating conditions described in (**j**). Error bars represent the SD from 4 biological replicates. p values were determined using Welch’s t-test. **l,** Alexa 647 MFI values from dead eGFP HEK293T transfected with or without GzmB gated prodrug mGELYTAC cocultured with NK92 cells (24 hrs). NK92 cells immunosuppressive and activating conditions described in (**j**). Error bars represent the SD from 4 biological replicates. p values were determined using Welch’s t-test.

Beyond TEV protease, we also tested selective gating by the protease granzyme B (GzmB). A protease of therapeutic interest^50,51^, GzmB is secreted by T cells and NK cells as part of the immune response to infection and cancer and contributes to autoimmune diseases^52^ (Fig. 3e). Thus, engineering GzmB-conditional therapeutics is an area of significant interest^53^. We generated an IGFPBP1-mGELYTAC construct similar to the above but with two GzmB cleavage motifs: one in the flexible disordered region of IGFBP1 and the other between IGFB1 and IGF2 (Fig. 3f, Extended Data Fig. 3b). We verified that this new mGELYTAC construct expresses and under the presence of recombinant GzmB is cleaved and mediates the internalization for Alexa-647 labeled rabbit IgG (Fig. 3g and Extended Data Fig. 3c). In practice an ungated mGELYTAC mediates the unrestricted IgG uptake in any cell type (Fig. 3h). However, for GzmB-gated mGELYTAC, since GzmB activity is confined in the immune synapse^54^, we hypothesize that its activity would likewise be confined only to NK cells and NK targeted cells (Fig. 3i). To test this, we cocultured eGFP HEK293T cells secreting GzmB-gated mGELYTAC or ungated mGELYTAC with NK92 cells either under immunosuppressive (TGF-β)^55^ conditions or immune-activating (IL2+ Il15+ PMA+ Ionomycin)^56, 57^ conditions. In non-targeted cells (live eGFP HEK293T), uptake was only mediated by ungated mGELYTAC regardless of activation state of the NK cells. No uptake was observed in either prodrug mGELYTAC condition (Fig. 3j). In NK cells, no IgG uptake was observed under immunosuppressive conditions but significant uptake of IgG in activated NK cells (Fig. 3k). Interestingly for ungated mGELYTACs, the NK cells exhibited a similar trend; this is likely due to the relative absence of IGF2R on the cell surface during immunosuppressive conditions and shuttling of IGF2R to the surface under activating conditions (Extended Data Fig. 3 d-e). Lastly, cells targeted by NK cells (dead HEK293T cells) exhibited a significant increase of IgG uptake during immuno-activating conditions over immunosuppressive conditions. Ungated mGELYTAC, on the other hand, mediated IgG uptake in putative target cells regardless of NK immune state (Fig. 3l).

#### CAR-T cell secreted GELYTACs

Another application of genetically encoded bifunctional molecules is secretion from CAR-T cells^58,59,60^. Like, other CAR-T secreted biologics^61,62^, GELYTACs initially had low secretion. To maximize their secretion, we optimized 3 aspects: (1) the signal peptide (BM40), (2) the order of GELYTAC and CAR virus transduction, and (3) a new mutation (R37A^63^) to prevent furin protease cleavage (Extended Data Fig. 4a-d). After optimization, we tested mCherry GELYTAC efficacy in a four-cell coculture model containing: primary T-cells expressing both α CD19 CAR and α mCherry GELYTAC or controls, CD19+ K562 cells, K562 cell secreting the mCherry target, and an eGFP+ K562 receiver cell. To compare the effects of CD19 CAR activation, we also included a 3-system coculture where the CD19+ K562 population was replaced with additional eGFP K562s. After confirming secretion of GELYTAC or controls (Fig. 4b, Extended Data Fig. 4e), we looked at mCherry uptake in receiver cells (Fig. 4c). At the 2-day mark and without CD19 cells, α mCherry GELYTAC secreting CAR-T cells mediated ∼10-fold higher mCherry uptake in receiver cells compared to that of untransduced primary T-cells. When CD19 cells were included, the mCherry fold change increased to ∼15 fold. Next, we analyzed the mCherry uptake in the CAR-T cell population (Fig. 4d). In inactivated CAR-T cells, we observed a modest uptake of around ∼2.6 fold over untransduced T-cells. However, once activated, that mCherry uptake increased up to ∼18 fold. In both the receiver and CAR-T analyses, the increase in mCherry uptake after CAR activation may be attributed to increased proliferation of the CAR-T cells after activation^64^ or elevated expression of IGF2R^65, 66^ (Extended Data Fig. 4f).

**Fig. 4:**
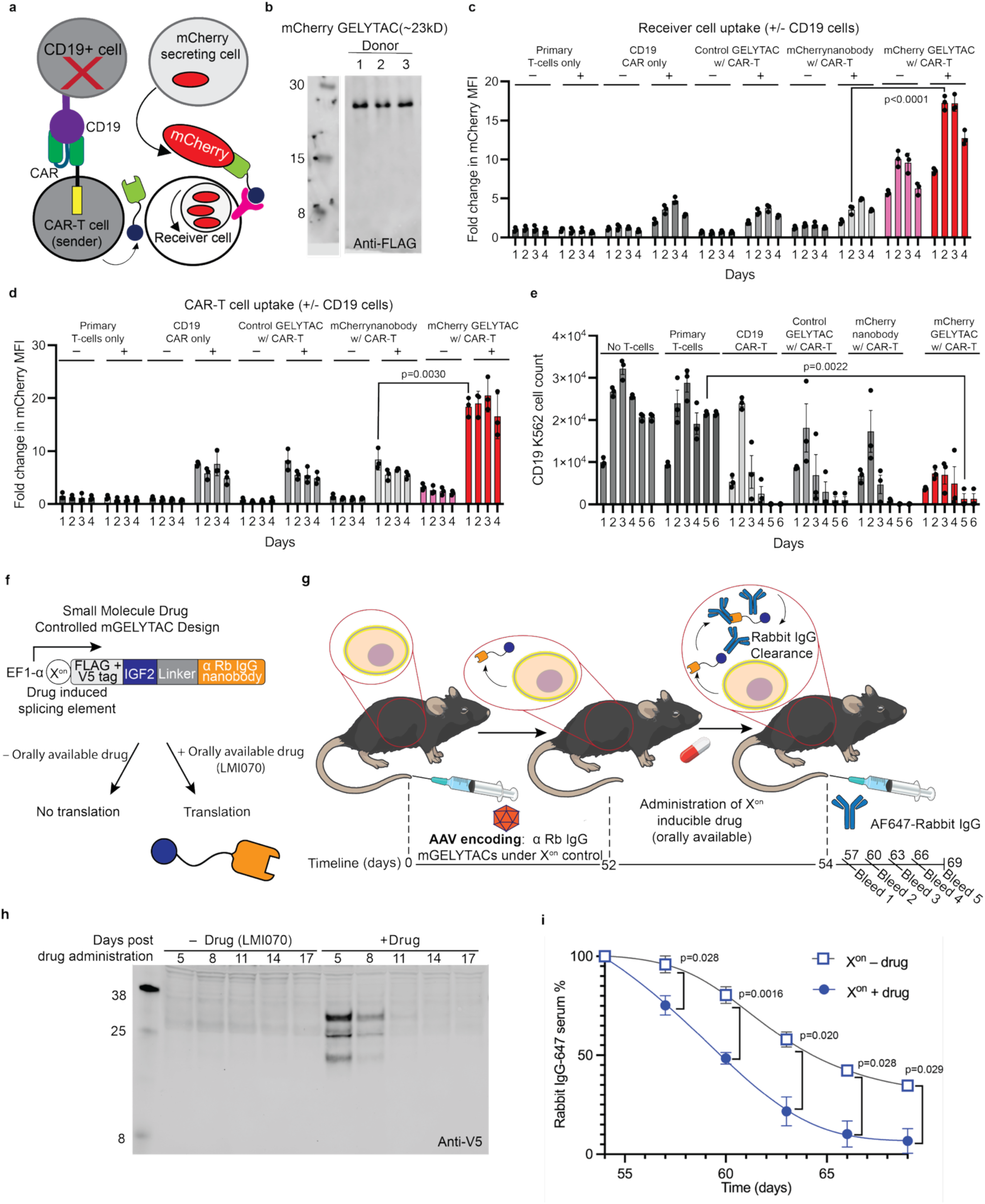
Cell-based synthetic biology applications of GELYTACs. **a**, Schematic depicting GELYTAC CAR-T proof of concept experiment. α CD19 CAR-T cells secreting α mCherry GELYTACs were cocultured with CD19+ K562s, mCherry secreting K562, and cytosolic eGFP K562 (receiver cells). **b**, α FLAG western blot of supernatant of CAR-T cells secreting mCherry GELYTACs from 3 donors. **c-d** Fold change in mCherry MFI (median fluorescence intensity) values from (**c**) receiver K562 or (**d**) CAR-T cells from the coculture experiment depicted in (**a**). Cells were cocultured for 1-4 days. Then, the cell mixture was analyzed using flow cytometry. Error bars represent the SD from 3 biological replications stemming from 3 different T-cell donors. p values were determined using Welch’s t-test. **e**, CD19 K562 cell counts after coculture with CAR-T GELYTAC cells. Cells were cocultured for 1-6 days, and an aliquot of cells was taken and counted with flow cytometry. Error bars represent the SD from 3 biological replications stemming from 3 different T-cell donors. **f**, Schematic of mGELYTACs under the control of drug induced splicing element (X^on^). **g**, Schematic of murine experiment for drug induced temporal selectivity of mGELYTACs. AAV encoding rabbit IgG targeting mGELYTACs under X^on^ was injected into C57BL/6 mice via tail vein injection. After 52 days, mice were orally administered LMI070. At day 54, mice were injected with Alexa-647 labeled rabbit IgG (20 mg/kg), and plasma was collected every 3 days post injection. Alexa-647 IgG levels were analyzed either via plate reader (**i**) or western blot (Extended Data Fig. 4d). **h**, α V5 western blot of murine plasma collected from C57BL/6 mice infected with AAVs encoding for α rabbit IgG mGELYTAC under X^on^ drug induced splicing element. Samples on the left are from a mouse treated with saline after AAV injection, and samples on the right are from a mouse treated with LMI070 after AAV injection. **i**, Alexa-647 rabbit IgG serum clearance as a function of time for mice treated with AAVs encoding for α rabbit IgG mGELYTAC under X^on^ control treated with saline (squares) or LMI070 (circles). Rabbit IgG clearance was measured via Alexa-647 plate reader assay on samples collected between 54-69 days post AAV injection. Error bars represent the SEM from 4 biological replicates for saline, and 3 biological replicates for LMI070. p values were determined using Welch’s t-test.

Finally, we assessed if arming CAR-T cells with GELYTACs impacts their ability to kill their target cells. We analyzed the cell count of CD19+ K562 cells co-cultured with CAR-T cells secreting mCherry GELYTAC or controls. Compared to T-cells transduced with just CD19 targeting CAR, we observe no decrease in killing ability (Fig. 4e). Altogether, our data suggests that GELYTACs could be integrated within CAR-T cells without comprising their cytotoxic capacity, while allowing for sustained target protein degradation.

#### Small molecule-inducible expression of GELYTACs

A concern for AAV based gene therapies is the constant expression of protein cargos. Thus, to afford temporal control of mGELYTACs at the level of transcription, we placed the transgene downstream of an X^on^ element, a small molecule-dependent alternative splicing module^44^. The X^on^ element is a modified *SF3B3* minigene that is prematurely terminated except in the presence of the small molecule splice modifier LMI070 (Branaplam) (Fig. 4f and Extended Data Fig. 4g). mGELYTAC constructs with (X^on^-mGELYTACs) or without the X^on^ element were transfected into HEK293T cells. As expected, only in the presence of LMI070 does X^on^-mGELYTAC express (Extended Data Fig. 4h, i).

We subsequently evaluated the performance of X^on^-mGELYTAC *in vivo*. An AAV8 construct encoding rabbit IgG-targeting X^on^-mGELYTAC (with a V5 epitope tag) was injected into the tail vein of C57BL/6 mice and LMI070 was orally administered 52 days later (Fig. 4g). Two days post-LMI070 treatment, Alexa-647-labeled rabbit IgG was injected into the tail vein, after which bleeds were performed every 3 days for 69 days. α V5 Western blots revealed that mGELYTACs were detectable only in plasma of LMI070-treated mice. Furthermore, circulating mGELYTAC was detected as early as 5 days after LMI070 administration with levels waning thereafter, (Fig. 4h) consistent with known drug pharmacology^44^. In addition to demonstrating LMI070-dependent expression, we also assessed mGELYTACs’ LMI070-dependent efficacy to deplete Alexa-647 labeled IgG. Samples taken every 3 days between day 54 and day 69 from LMI070-treated mice exhibited significant decreases in Alexa-647 signal across all timepoints compared to untreated mice (Fig. 4i). These data were corroborated by western blot analysis of the IgG heavy and light chains (Extended Data Fig. 4j).

### Clearing Endogenous IgGs with gene therapy-delivered mGELYTAC

Having shown effective clearance of exogenous rabbit IgG with mGELYTAC (Fig. 2), we next explored whether *in vivo* mGELYTAC production could lower steady-state levels of endogenous murine IgG, a therapeutic goal of many autoimmune disease treatment paradigms. In C57BL/6 mice, IgGs consist of 2 light chains (κ or λ) and 2 heavy chains (1, 2b, 2c or 3), with individual chains encoded by independent genes (Fig. 5a). Different chains combine to form various IgG isotypes, each with their specific function and role in autoimmune diseases^67^. Thus, targeting specific IgG subtypes is desirable^68^. We reasoned that mGELYTACs comprising isotype-specific nanobodies could achieve this goal^32^.

**Fig. 5:**
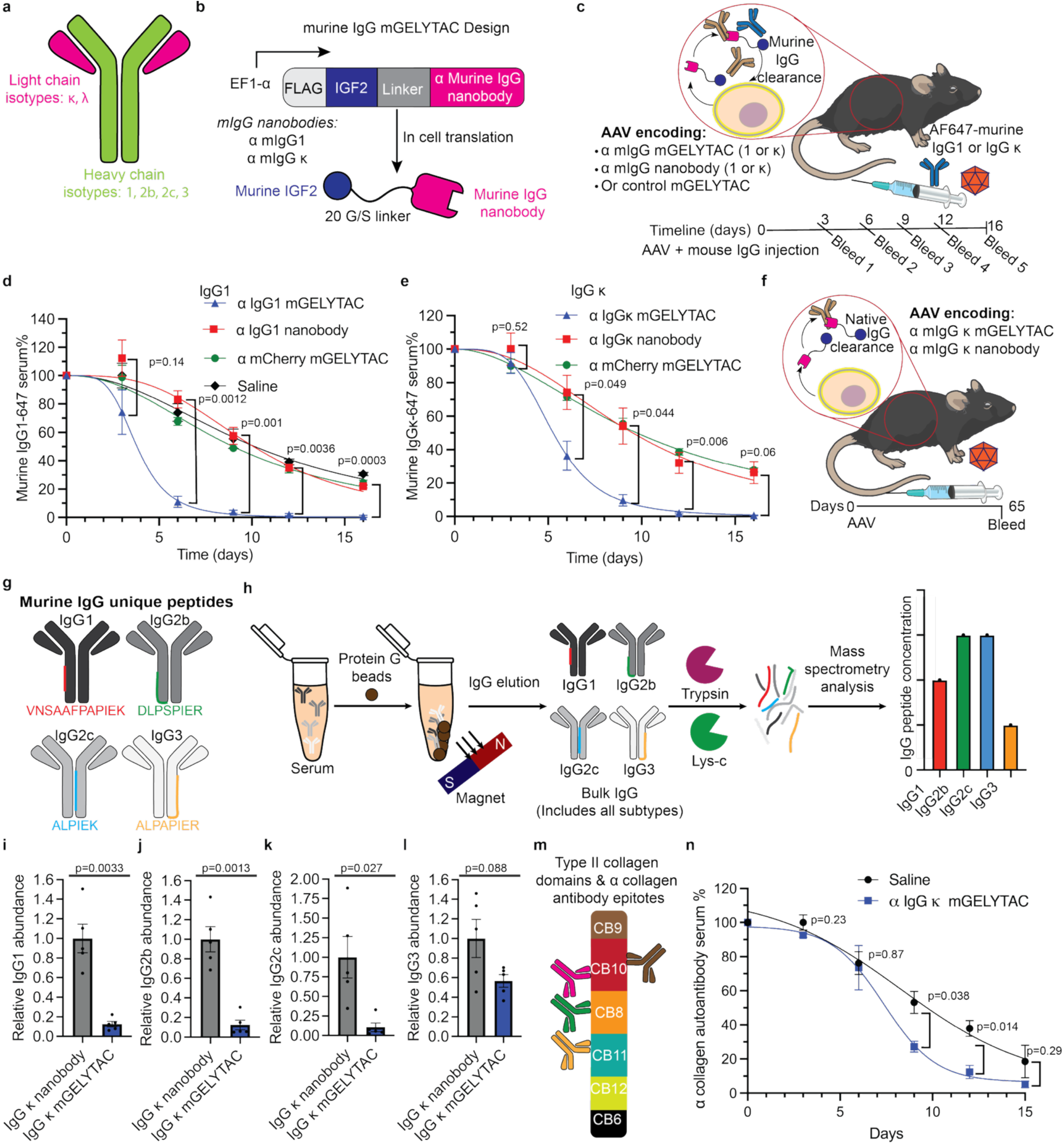
Gene therapy delivered mGELYTACs clear endogenous murine IgG. **a**, Schematic depicting murine IgG types. **b**, Schematic of murine IgG targeting mGELYTACs. Murine IgG targeting mGELYTACs contain a nanobody targeting either IgG1 or IgG κ. **c**, Schematic outlining murine-IgG clearance in vivo via measuring an exogenous Alexa-647 labeled IgG. Alexa-647 labeled murine IgG was injected into C57BL/6 mice at a level subceeding endogenous amounts (1/165^th^ for IgG1 or 1/45^th^ for IgG κ) along with AAV8 encoding for murine IgG targeting mGELYTAC, or murine IgG nanobody, or control mGELYTAC (targeting mCherry^59^). **d**, Plate reader analysis of murine Alexa-647 IgG1 serum clearance as a function of time for mice treated with AAVs encoding for murine IgG1 mGELYTAC (triangles), murine IgG1 nanobody (squares), control mGELYTACs (targeting mCherry, circles), or saline (rhombuses). Error bars represent the SEM from 3 biological replicates for murine IgG1 mGELYTACs, 3 biological replicates for murine IgG1 IgG nanobody, 3 biological replicates for control (mCherry) mGELYTACs, and 3 biological replicates for saline. p values were determined using Welch’s t-test. **e**, Plate reader analysis of murine Alexa-647 IgG κ serum clearance as a function of time for mice treated with AAVs encoding for murine IgG κ mGELYTAC (triangles), murine IgG κ nanobody (squares), or control mGELYTACs (targeting mCherry, circles). Error bars represent the SEM from 3 biological replicates for murine IgG κ mGELYTACs, 3 biological replicates for murine IgG κ IgG nanobody, and 3 biological replicates for control (mCherry) mGELYTACs. p values were determined using Welch’s t-test. **f**, Schematic of murine experiment for endogenous IgG clearance. **g**, Schematic of unique peptides on IgG heavy chain subtypes present in C57BL/6 mice that could be used to quantify using bottom-up proteomics. **h**, Schematic of bottom-up proteomics assay to quantify IgG heavy chain subtypes in C57BL/6 mice. IgG was first pulled down and subsequently digested into peptides. Unique peptides specific to IgG heavy subclasses were then quantified using mass spectrometry. **i-h,** IgG subtype plasma relative abundance from C57BL/6 mice treated 65 days prior with AAV8 encoding IgG κ mGELYTACs or nanobody determined from mass spectrometry. Error bars represent the SEM from 5 biological replicates. p values were determined using Welch’s t-test. **i,** IgG1 **j,** IgG2b **k, l,** IgG3 **m**, Schematic showing the collagen domains that disease-causing α collagen antibodies bind to. **n**, Murine α collagen antibody cocktail-647 (100 mg/kg) serum clearance as a function of time for mice treated with AAVs encoding for IgG κ (squares). AAV was injected seven days prior to cocktail administration. α Collagen murine IgG clearance is measured via Alexa-647 plate reader assay on samples collected between 3-15 days post IgG injection. Error bars represent the SEM from 3 biological replicates. p values were determined using Welch’s t-test.

We designed two mGELYTAC constructs against IgG1 and IgG κ, respectively (Fig. 5b). The IgG1 and IgG κ mGELYTACs showed good expression in cell culture and maintained their selectivity for their respective IgG targets (Extended Data Fig. 5a-d). Specifically, IgG1 mGELYTAC binds IgG1 subtypes containing either κ or λ light chains. IgG κ mGELYTAC targets all IgG κ subtypes containing either IgG1, IgG2a, IgG2b, or IgG3 heavy chains. In a coculture experiment with eGFP HEK293T sender cells with X16C.8 lymphocyte receiver cells, we showed uptake of Alexa-647-labeled murine IgG by both mGELYTACs with the expected IgG subtype specificities (Extended Data Fig. 5e,f). To confirm degradation of internalized IgG, we analyzed the conditioned media as well as lysates of the receiver cells by Western blot (Extended Data Fig. 5g,h)^31^.

We next confirmed that the IgG-specific FLAG-tagged mGELYTACs expressed *in vivo* upon viral transduction. We infected C57BL/6 mice by tail-vein injection of either mGELYTAC in an AAV8 vector (Extended Data Fig. 5i). Plasma was collected every three days post-injection for 15 days and mGELYTAC expression was confirmed by α FLAG Western blot (Extended Data Fig. 5j,k). We also validated the ability of circulating mGELYTACs to clear IgG, first using exogenously administered Alexa-647 tagged IgG1 or IgG κ. We injected these proteins into mice at low levels compared to known concentrations of endogenous murine IgGs (1:165 for IgG1 and 1:46 for IgG κ)^40^, along with the respective AAV8-mGELYTAC viral particles (Fig. 5c). Plasma samples were collected every 3 days for 16 days and analyzed on a fluorescence plate reader. When compared to controls, both IgG1-targeting and IgG κ-targeting mGELYTACs significantly accelerated the clearance rate of their respective IgG isotypes by ∼2.5 fold and ∼2 fold, respectively (Fig. 5d,e).

Ultimately, the therapeutic potential of mGELYTACs hinges on their ability to reduce endogenous IgG levels. To this end, we injected AAV8 encoding for IgG κ targeting mGELYTAC in mice (Fig. 5f) and subsequently utilized liquid chromatography-tandem mass spectrometry (LC-MS/MS) to quantify serum levels of IgG ^70, 71^. Murine IgG heavy chains contain subtype-specific regions that can be detected and quantified by bottom-up mass spectrometry proteomics (Fig. 5g). To this end, we quantified murine IgG isotype levels by purifying bulk murine IgG via protein G affinity capture and subsequent tryptic digestion to generate peptides that were quantified by LC-MS/MS (Fig. 5h). We analyzed plasma from mice 65 days post AAV8 treatment and found robust expression of IgG κ mGELYTACs and IgG κ targeting nanobody (Extended Data Fig. 5l). Accordingly, we observed a ∼87% drop in IgG1 (Fig. 5i), ∼87% drop in IgG2b (Fig 5j), ∼89% drop in IgG2c (Fig. 5k), and an apparent ∼45% drop in IgG3 (Fig. 5l) in mice treated with AAV8-IgG κ-GELYTAC compared to mice treated with AAV8-IgG κ-specific nanobody. In parallel, we verified the reduction of IgG2b using an ELISA (Extended Data Fig. 5 m-n) and were able to see a similar depletion pattern.

Lastly, we tested mGELYTAC’s potential to reduce disease phenotypes in a rheumatoid arthritis disease model^72^. In this model, a cocktail of mouse collagen targeting antibodies (Fig. 5m) are passively transferred to mice. These antibodies bind to collagen on the joints, mediating inflammation via the recruitment of the innate immune system causing joint swelling and damage ^72^. First, we showed that mGELYTACs were able to mediate the uptake and clearance of these disease-causing antibodies in cell culture and *in vivo* (Fig. 5n). Then, we injected these antibodies followed by AAV8 encoding for IgG κ mGELYTAC into mice and assessed the disease phenotype after 13 days (Extended data Fig. 5p). A trend of reduced swelling in mice treated with AAV encoding mGELYTAC compared to AAV encoding for IgG κ targeting nanobody was observed, which is consistent with Efgartigimod alfa (Extended data Fig. 5q). The disease phenotype was judged by two blinded individuals assessing the degree of inflammation (see methods). Mice treated with AAV encoding for mGELYTAC exhibited significantly lower disease score compared to mice treated with the AAV encoding for nanobody (Extended Data Fig. 5r) As an alternative readout of disease, we also utilized a rotarod test for these mice^73^. After 8 days post collagen antibody administration, we observed that mice treated with AAV8 encoding for mGELYTAC stayed on the rod significantly longer than mice treated with AAV encoding nanobody control (Extended Data Fig. 5s) All together, these results suggest that mGELYTACs can have therapeutic efficacy.

## Discussion

Extracellular targeted protein degradation is an emerging therapeutic modality that, to date, has centered on exogenously administered biologics and bioconjugates. The GELYTACs reported here are delivered by gene therapy vectors for continuous *in vivo* production after a single administration. We demonstrated proof-of-principle by developing a GELYTAC that depletes IgG molecules for at least two months following a single dose. This stands in contrast to other IgG depleting therapeutics, such as Biohaven’s GalNAc-peptide conjugate BHV-1300 which require weekly dosing^30^. To circumvent low serum half-life, others have developed eTPD modalities that are catalytic ^28, 74^ or contain conditionally active domains^75^. Although catalytic extracellular degraders reportedly provide an increased efficacy window, target (IgE) levels begin to rebound in mice after 14 days and return to serum levels around 21 days^28^. Circulating IgG levels can be also reduced by B cell depletion using the CD20 antibody Rituximab or by emerging B cell-targeted CAR-T therapies^76^. These approaches are insufficient for some patients because there exists a CD20 negative B-cell population that produces autoantibodies^77^. Also, CAR-T therapeutics for autoimmune disease have significant side-effect profiles^78^. Thus, a gene therapy GELYTAC-based strategy for IgG depletion may have a unique position in the development of autoimmune disease therapeutic development.

The GELYTAC strategy is also amenable to a variety of spatial and temporal control mechanisms on either the protein or genetic level. In our study we provided a protein level example with protease-gated receptor engagement. In the future, similar modalities could potentially modulate the depletion of proteins only in disease sites with higher levels of proteases such as Granzyme B in autoimmune disease or matrix metalloproteases in cancer. We also included a genetic example selectivity using a small molecule to gate alternative splicing. In theory most genetic strategies to enhance selectivity would be applicable to GELYTACs. Examples include AAV serotypes that target specific disease states^79^, conditional antigen gating^80^, and protease gated secretion^81^.

While we also studied the safety profile of mGELYTACs in mice in the context of ADAs and IGF2 clearance, IGF2 also binds IGF1R and the insulin receptor (IR) in addition to IGF2R and can signal through them. To advance GELYTACs therapeutically, growth signaling and long-term tolerability will need dedicated studies. This is potentially addressable by protein engineering to generate IGF2 variants blinded to IGF1R and IR binding.

Gene therapy delivered mGELYTAC, like other genetic medicines, potentially faces increased regulatory hurdles due both viral and biologic safety considerations^82^. Nevertheless, there has been considerable progress in the field of viral delivery vehicles, paving the way for the future^83^. As we have shown with our proof of principle with GELYTAC secreting CAR-T cells, GELYTACs have additional applications beyond gene therapy, and could be compatible to pair with in vivo delivered CAR T platforms. In addition to what we have shown in this study, GELYTACs can have potentially broad application being integrated within different synthetic biology paradigms or with alternative delivery vehicles such as mRNA-LNPs and VLPs^84^. These improvements include making delivery vehicles that are less immunogenic^85^, easier to manufacture^86^, and more selective^87^.

## Methods

### I. Construct cloning

AAV mGELYTAC plasmids were generated using either pAAV-EF1a or pAAV-hSyn backbones. To enhance protein expression, all constructs included a 3′ woodchuck hepatitis virus post-transcriptional regulatory element (WPRE). Constructs also included the inverted terminal repeat (ITR) regions to enable AAV packaging. For drug induced splicing versions, the X^on^ element was positioned upstream of mGELYTAC sequences. CAR-T and GELYTAC retroviral constructs were cloned into the MSGV1 vector.

To generate new plasmids designs, the gene insert(s) were either generated with PCR (Q5 polymerase master mix, New England Biosciences) or ordered from Twist Biosciences (codon optimized for murine). The overhangs on both ends of the inserts were designed to have an annealing temperature of ∼60°C-70°C with the pAAV backbone. The digested backbones were generated with BamHI (New England Biosciences) and EcoRI (New England Biosciences). The full gene cassettes were ligated using HiFi master mix (1 hr reaction at 50°C). Restriction enzymes, Q5 mastermix, and Hi-Fi master mix were implemented based on manufacturer instructions. Plasmids were then transformed into stable competent *E. coli* (New England Biosciences) and then plated on an LB agar plate with ampicillin. Colonies were then screened by sequencing (Plasmidsaurus) to identify plasmids with the correct sequence.

### II. Cell culture

Immortal cell lines: all cell lines used were less than passage 25. HEK293T cells (ATCC), MC38 (Sigma Aldrich), J77A.1 (ATCC), AML12 (ATCC), and BV2 were cultured in a DMEM (Gibco) supplemented with 10% fetal bovine serum (FBS), 1% Glutamax (Gibco), 100 units/mL penicillin, and 100 mg/mL streptomycin at 37 °C under 5% CO2. BMA3.1 (ABM), X16C8.5 (ATCC), K562s (ATCC) was cultured in in RPMI (Sigma Aldrich) supplemented with 10% FBS, 1% Glutamax (Gibco), 100 units/mL penicillin, and 100 mg/mL streptomycin at 37 °C under 5% CO2. X16C8.5 cells were supplemented with an additional 0.05 mM 2-mercaptoethanol. NK92 cells (ATCC) were cultured in MyeloCult H5100 medium (StemCell Technologies) supplemented with 100 units/mL penicillin, and 100 mg/mL streptomycin, 6.3% horse serum, and 10 μg/500mL IL-2.

Coculture protocols: On day-1, 6e5 eGFP HEK29T were plated on human fibronectin (HFN) coated 6-well plates. On day 0, eGFP HEK293T cells were transfected with 2000 ng of plasmid DNA encoding for mGELYTACs or controls. DNA was added to 200 μL of blank DMEM followed by 10 μL of polyethylene imine (PEI) (1 mg/mL, Polysciences, 24765). Transfection mix was allowed to incubate for 20 min before adding to eGFP cells dropwise. On day 1, transfected eGFP cells were lifted with trypsin. 1.5e5 (0.5 mL) eGFP HEK293T cells secreting mGELYTAC or controls were mixed with 1.5e5 (0.5mL) murine receiver cells. Coculture was supplemented with 100nM of Alexa-647 labeled target antibody and plated on HFN coated 12 well plates. Cells were analyzed on day 2 using flow cytometry.

### III. Flow cytometry

300 μL of X16C8.1 cell culture or 300 μL of adherent cell culture (lifted with enzyme free cell dissociation buffer) was transferred to 96-well V-bottom plate and spun down (500g for 1 min). Cells were then washed 1x PBS w/ 0.5% supplemented bovine serum albumin (BSA), EDTA (2.5 mM, and 2 units of DNAse I (Thermo Fisher AM2238)/ mL). Cells were then incubated with SYTOX Blue (1:1000 dilution) for 5 min. Flow cytometry was performed with MACSQuant, BioRad ZE5, or Agilent Quanteon flow cytometers, and analysis was performed using the FlowJo software package. Gating was performed on single cells and live cells (Supplemental Fig. 1). This instrument is equipped with a 405-nm violet laser, a 488-nm blue laser, a 561-nm green laser and a 639-nm red laser.

### IV. Cell culture labeling for confocal imaging

Coverslips were sterilized under a UV light for 10 min before they were coated with HFN. The specific coculture conditions are listed in the coculture section, but with half of the indicated values to adjust for the 24-well plate format. Post-coculture, cells were carefully rinsed 3x with PBS and treated with 4% paraformaldehyde (PFA) in PBS for 15 min. Cells cultured on coverslips in 24-well plates were rinsed three times with room temperature (RT) 1x PBS to remove residual fixative. Cells were incubated with 1x PBS containing 3% normal bovine serum (NBS) and 0.03% Triton X-100 for 30 minutes at RT with moderate shaking. Primary antibodies were diluted in 1x PBS with 3% normal bovine serum (NBS), and the blocking solution was replaced with the primary antibody solution; a secondary-only control (no primary antibody) was included. Cells were incubated with the primary antibody overnight at 4 °C with gentle shaking or tilting. The following day, cells were washed three times with 1x PBS for 5 minutes each at RT. Secondary antibodies were diluted (typically 1:500) in 1x PBS containing 3% NBS and applied to cells for 3 hours at RT with gentle shaking, protected from light thereafter. After staining, cells were washed three times with 1x PBS for 5 minutes each at RT. Coverslips were then mounted onto glass slides using mounting media prior to imaging. Confocal images were captured on a Leica STELLARIS confocal microscope using a Leica HC PL APO CS2 40X oil objective, with a 405 nm diode and a white-light laser configured at specific dye excitation wavelengths.

### V. AAV Virus Generation

All adeno-associated viral vectors were produced by the Stanford University Gene Vector and Virus Core (RRID:SCR_023250). In short, AAV-293 cells (Agilent Technologies) were triple transfected. The supernatant was then subjected iodixanol gradient ultracentrifugation, buffer exchange, and ultrafiltration. Genomic titers were calculated via digital quantitative PCR (QIAcuity One, QIAGEN, Germantown, MD) using primers/probes targeted to WPRE and ITR sequences.

### VI. Animal Care

All experimental procedures adhered to NIH regulations and were approved by the Stanford University Administrative Panel on Laboratory Animal Care. For all animal experiments, male C57BL/6 mice were used.

For exogenously supplied target animal experiments C57BL/6 mice were injected via tail vein with 1e11 viral genomes of AAV expressing mGELYTAC constructs, and 12.5 mg/kg – 40 mg/kg fluorescently labeled IgG. Blood samples from cheek bleeds were then taken 3, 6, 9, 12, and 15 days post injection. Blood samples were centrifuged 500 xg for 5 mins and only plasma was collected for further experiments. For mouse IgG fluorescence experiments C57BL/6 mice were injected via tail vein with 1e11 viral genomes of AAV expressing mGELYTAC constructs and either 0.25 mg/kg fluorescently labeled murine IgG 1λ-AF647, or 5 mg/kg fluorescently labeled murine IgG2aκ. For the experiment outlined in Extended Data Fig. 6b, AAV was injected 1 week prior to administration of 75 mg/kg murine α collagen IgG-647. Blood samples from cheek bleeds were taken 3, 6, 9, 12, and 15/16 days post injection and centrifuged as before to isolate plasma.

For endogenous IgG clearance experiments. Animals were injected with 1e11 viral genomes, and serum collected at day 65 was used for analysis. For the rheumatoid arthritis disease model, mice were first injected intraperitoneal with 200 mg/kg α collagen IgG cocktail. 3 days later mice were injected with 100 μg LPS to induce inflammation.

All animals were weighed pre– and post-injection to monitor weight fluctuations due to agent administration. No abnormal changes were observed, and animals were fully mobile and alert. Animals were kept on a standard lab diet and water at all times.

### VII. Plasma collection

Plasma was collected from mice every three days through submandibular bleeding using fine needles (BD) into lithium heparin tubes (Sarstedt, 20.1282.100), and samples were immediately placed on ice. Blood was then centrifuged at 5,000 rpm for 5 minutes at 4 °C. The plasma (top layer) was carefully aliquoted and stored at −80 °C. This procedure was based on a previously established protocol^88^.

### VIII. Recombinant protein expression

Recombinant mGELYTAC (Supplemental Fig. 2 a,b) and IGF2-mouse serum albumin (MSA) (Supplemental Fig. 2c) were produced using an Expi293F expression system (Thermo Fisher A14635). Both protein constructs were cloned into a pcDNA3.4 expression vector. Expi293F cells were transfected with Expifectamine (Thermo Fisher A14524) according to manufacturer instructions. 5 days post transfection, cells were spun down at 1000xg for 15 min. The supernatant was then filtered using a 10 μm filter and before flowing through a Ni-NTA column (Cytvia 17524801, using the sample pump on Cytiva AKTA Pure). The samples were then washed with 10 column volumes of Tris-HCl pH8 buffer and eluted with 300mM Imidazole (in Tris-HCl pH8 buffer). The Ni-NTA eluent was then further purified using size exclusion chromatography (Cytiva 28989336). Protein concentrations were quantified using a Thermo Fisher Nanodrop.

### IX. LYTAC generation

LYTAC generation was based on previously published methods^25^. Goat α rabbit IgG (Jackson Immunoresearch 111-005-144, dialyzed into PBS) was conjugated to NHS-DBCO at room temperature for 8 hrs (25 equivalents). After another round of dialysis into fresh PBS, IgG-DBCO conjugates were conjugated to Tri-GalNAc azide (Supplemental Fig. 2d) Tocris 8015, 25 equivalents) at room temperature for 24 hrs. After the reaction, the mixture was then dialyzed again into fresh PBS to yield the final LYTAC (Supplemental Fig. 2e).

### X. Western blot protocols

Conditioned media: conditioned media from transfected HEK293T cells was collected (cells were removed by centrifugation) and LDS Sample Buffer (4X) (for reduced blot 4x loading buffer was supplemented with 10% 1M DTT) and boiled at 95 °C for 10 min. The recommended volume (15 μL for 26-well gel or 30 μL for 18-well gel) was loaded on an SDS–PAGE (10% Bis-Tris gel), then transferred to a nitrocellulose membrane. After transfer, the blot was blocked with Odyssey Blocking Buffer (PBS or TBS) (LI-COR) for 1 h at room temperature with gentle shaking. Membranes were stained with M2 α FLAG (Sigma Aldrich) for 1 hr at room temperature or at 4 °C with gentle shaking, then washed three times with TBS-T for five minutes each. The membrane was then incubated with 800CW goat α mouse IgG (1:10,000) in Odyssey Blocking Buffer (PBS or TBS) for 1 h at room temperature with gentle shaking. Membranes were washed three times with TBS-T, then imaged using an OdysseyCLx imager (LI-COR). Quantification of band intensities was performed using Image Studio Software (LI-COR).

Murine plasma blots: for plasma blots 6 μL of murine plasma was diluted in 39 μL PBS. Diluted plasma samples were then mixed with 15 μL 4x LDS sample buffer (Thermo Fisher, NP0007) or 4x loading buffer (LiCor Bio, 928-40004). Samples were mixed and boiled for 10 min at 95 °C before loading into a 10% Bis-Tris gel and ran at 140 V for 45 min. The gel was then transferred into a nitrocellulose membrane and blocked with Intercept PBS Blocking Buffer (LI-COR Bio, 927-70001) for 1 h at room temperature with gentle shaking. Membranes were then incubated with chicken α FLAG (Abcam, AB1170) or chicken α V5 (Thermo Fisher, A190-118A) antibody overnight at 4 °C with gentle shaking. The following morning membranes were washed three times with TBS-T for 5 min with gentle shaking. Washed membranes were then incubated with IRDye 800CW Donkey α Chicken secondary antibody (1:10,000) (LI-COR Bio, 926-32218) in Intercept PBS Blocking Buffer for 1 hr at room temperature with gentle shaking. Membranes were washed three times with PBS-T for 5 min and imaged using an OdysseyCLxImager (LI-COR). Quantification of band intensities was performed using Image Studio Software (LI-COR).

Dot blot: 0.5 ug of murine antibody subtypes were added onto nitrocellulose and allowed to air dry for 10 minutes. After drying, the nitrocellulose membrane was blocked using Odyssey Blocking Buffer (PBS) for 1 hr). Then the membrane was incubated with conditioned media from mGELYTAC transfected HEK293T cells (3 days post transfection) for 1 hr. The blot was then washed three times with TBS-T for 5 min. The blot was then incubated with rabbit-α FLAG antibody (Proteintech 20543-1-AP) (1:1000) for 1 hr and then washed with the wash conditions listed above. Finally, the blot was stained with goat-anti Rabbit IR800 secondary antibody for 30 min, and washed a final time, again with the wash conditions stated above. The blot was imaged using an OdysseyCLxImager (LI-COR).

### XI. Plate reader assay

20-100 μL Conditioned media or murine plasma (neat plasma or plasma, diluted in PBS) were plated in a clear bottom/black 96-well plate (Corning Costar, 3603) and analyzed using a Tecan M1000/Pro for Alexa-647 signal (excitation: 650 nm, emission: 671 nm). In downstream data analysis plots (Fig 2E, 2H, 4D, 5D, 5E), the Y-axis 100% represents the day 3 saline or nanobody only control (when no saline control present) Alexa-647 reading. “Day 0” reading is carried over from day three saline or nanobody only control (when no saline control present) treatment Alexa-647 reading.

### XII. Autoantibody ELISA

Rabbit IgG isotype control (Thermo Fisher) was labeled with EZ-Link NHS-LC-LC-Biotin (Thermo Fisher, 21343) dialyzed and purified using size exclusion chromatography on fast protein liquid chromatography (FPLC) to yield monomeric biotinylated rabbit IgG. The reaction conditions for NHS bioconjugation were 50 μM NHS biotin in the antibody (5 mg/mL) storage buffer. Streptavidin 96-well plates (Thermo Fisher, 15120) were coated with biotinylated rabbit IgG (0.5 mg/mL) in a washing buffer (Thermo Fisher, 37535 supplemented with 0.05% Tween20) for 2 hrs. After 3 200 μL washes, diluted plasma (10%, 90% wash buffer) was incubated for 2 hrs. For the α FLAG positive control, 20 μL of murine plasma was incubated with 2 μL of M2 murine α FLAG for 1 hr at room temperature. Following plasma incubation, the plate was washed 3 times with wash buffer and stained with chicken anti murine HRP (1:1000) (Thermo Fisher, A15975). After a 30 min incubation, the plate was washed 3 time and incubated with HRP substrate (Thermo Fisher, 34028) for 15 minutes before reading on a plate reader for absorbance (450 nm).

### XIII. Source of primary human T cells (based on previous published methods^89^)

Buffy coats from healthy donors were purchased from the Stanford Blood Center under an IRB-exempt-protocol. Leukopaks from healthy donors were purchased from StemCell Technologies. Primary human T cells were purified by negative selection using the RosetteSep Human T cell Enrichment kit (StemCell Technologies) and SepMate-50 tubes. T cells were cryopreserved at 2 × 10^7^ cells per ml in CryoStor CS10 cryopreservation medium (StemCell Technologies) until use.

### XIV. Retrovirus production (based on previous published methods^89^)

Retroviral supernatant was packaged using 293GP cells and the RD114 envelope plasmid. In brief, 11 μg RD114 and 22 μg of the corresponding MSGV1 transfer plasmid were delivered to 293GP cells grown on HFN coated 150 mm dishes (Corning) to 80% confluency by transient transfection with Lipofectamine 2000 (Thermo Fisher Scientific). The medium was replenished every 24 hr. The retroviral supernatant was collected 48 and 72 h after transfection. The supernatants from replicate dishes were pooled, centrifuged to deplete cell debris and stored at −80 °C until use.

### XV. CAR-T cell generation (based on previous published methods^89^)

At day 0, primary human T cells were thawed and activated with anti-CD3/CD28 human T-Expander Dynabeads (Thermo Fisher Scientific) at a 1:1 bead to cell ratio. On day 2, virus-coated culture plates were prepared on non-tissue-culture-treated 12-well plates that had been precoated with RetroNectin (Takara Bio) according to the manufacturer’s instructions, by incubating with 1 ml of retroviral (encoding for GELYTAC) supernatant (2 × 10^7^–5 × 10^7^ TU ml^−1^) and centrifugation at 3,200 rpm and 32 °C for 2 h. The supernatant was subsequently aspirated off of the wells and 0.5 × 10^6^ T cells were added in 1 ml of T cell medium comprising RPMI1640 (Thermo Fisher Scientific), 10% FBS, 100 U ml^−1^ penicillin (Gibco), 100 mg ml^−1^ streptomycin (Gibco), 2 mM L-glutamine (Gibco), 10 mM HEPES (Gibco) and 40 U ml^−1^ rhIL-2 (Peprotech). After addition of the T cells, the plates were gently centrifuged at 1,000 rpm for 2 min then incubated for 24 h at 37 °C under 5% CO_2_. On day 3, this process was repeated but with retrovirus encoding for CD19 CAR. Dynabeads were removed on day 4 by magnetic separation. Cells were maintained between 0.4 × 10^6^ and 2 × 10^6^ cells per ml and expanded until day 10–12.

### XVI. CAR-T coculture

mCherry uptake assay: In the CD19+ coculture, 5e4 CD19+ K562 (with eGFP and eBFP), 5e4 Igκ mCherry K562, 5e4 eGFP K562, and 2e5 GELYTAC secreting CAR-T cells (or control secreting CAR-T cells) were cocultured in 2mL RPMI supplemented with IL2. In the CD19+ coculture, 5e4 Igκ mCherry K562, 1e5 eGFP K562, and 2e5 GELYTAC secreting CAR-T cells (or control secreting CAR-T cells) were cocultured in 2mL RPMI supplemented with IL2.

Cell killing assay: 1.25e5 CD19+ K562s were cocultured with 2.2e5 GELYTAC secreting CAR-T cells in 3mL RPMI supplemented with IL2.

### XVII. IgG ELISA

Murine IgG ELISA (IgG2b: Thermo Fisher, 88-50430, IgG2a: 88-5042088) was conducted to assess IgG levels in murine plasma following the manufacturer’s protocol. Briefly, plates (Thermo Fisher, 439454) were coated overnight at 4C with purified α murine IgG capture antibody. The next morning, plates were washed twice with washing buffer, 1x PBS containing 0.05% Tween-20. After washes, plates were blocked for 2 hours at room temperature with blocking buffer supplied in the kit. Murine IgG standard included in the kit was prepared according to the kit protocol and murine plasma samples were diluted 1:20,000 in 1x Assay Buffer A. Standard samples and plasma samples were added to the plates and incubated with HRP-conjugated α mouse IgG polyclonal antibody for 3 hours at room temperature while shaking at 400rpm. Subsequently, the plates were washed four times with washing buffer described previously and incubated with TMB substrate solution for 15 minutes. Lastly, stop solution consisting of 2N Sulfuric acid was added and plates were read using a microplate reader set to read absorbance at 450nm. A curve was generated from the standard samples and used to calculate concentrations for the plasma samples.

### XVIII. Luciferase Assays

For nano luciferase assays: 100 μL of 0.165 mg/ml furimazine in (45 % PBS, 23 % ethanol, 32 % polyethylene glycol 300) was injected via retroorbital injection. After 10 minutes, the organs were dissected and plated for analysis on an IVIS Lumina Series III (Perkin Elmer) imaging system with 30-second exposures and medium binning. Images were analyzed and normalized using Revvity LivingImage software.

For HiBit/LgBit assays: 50 μL of LgBit (32mg/mL) and 50 μL of 0.165 mg/ml furimazine in (45 % PBS, 23 % ethanol, 32 % polyethylene glycol 300) were injected via 2 separate retroorbital injections into different fornix conjunctival membranes. After 10 minutes, the organs were dissected and plated for analysis on an IVIS Lumina Series III (Perkin Elmer) imaging system with 5-minute exposures and medium binning. Images were analyzed and normalized using Revvity LivingImage software. The mouse organs in Extended Fig. 2c and Extended Fig. 2e are the same mouse but taken using the fluorescence setting (Extended Fig. 2c) or the bioluminescence setting (Extended Fig. 2e).

### XIX. Oral drug admin

In drug induced splicing experiments, a single 50 mg/kg dose of LMI070 (MedChemExpress, HY-19620) or vehicle was administered by oral gavage 4 weeks after AAV intravenous injection.

### XX. Prodrug assays

#### TEV protease

TEV protease prodrug assays follow the standard coculture assay up until day 3, with exception that 185 nM of Alexa-647-labeled rabbit IgG was used. On day 3, 500nM TEV protease (UC Berkeley QB3 MacroLab) was added. On day 4 cells were analyzed via flow cytometry.

#### Granzyme B

On day 0, eGFP HEK293T cells were seeded in T25 flaks (5 mL of 3.5e5 cells/mL). On day 1, transfection mixes for each well were made with 500 μL of blank DMEM mixed with 5000 ng of DNA followed by 25 μL of PEI (1mg/mL). On day 2, granzyme B (375 nM) and Alexa-647-labeled rabbit IgG (200 nM) were added. On day 3, the HEK293T conditioned supernatant was transferred to MC38 cells (seeded 1 day prior at 3e5/mL in 12 well plate and stably expressing serpinb9-P2A-mCherry). On day 4, MC38 cells were lifted and analyzed via flow cytometry.

### NK cell prodrug assay

#### eGFP HEK293T cells

On day 0, eGFP HEK293T cells were seeded in 12-well plates (1 mL of 3.5e5 cells/ mL). On day 1, transfection mixes for each well were made with 100 μL of blank DMEM mixed with 1000 ng of DNA followed by 5 μL of PEI (1mg/ mL). The transfection mix incubated for 20 min before addition to eGFP HEK293T cells.

#### NK cell culture

For immunoactiving conditions: On day 1: 5ug of IL2 and 10ug of IL15 were added to 16mL of maintenance NK92 cell culture. For immunosuppressive conditions, maintenance media containing IL2 from 16 mL of NK92 cell culture was exchanged with 16mL RPMI containing 20 ug of TGF-β. On day 2 before adding NK cells to HEK293T cells, ionomycin and PMA were added to the immunoactivting conditions <u>only.</u>

#### Coculture

On day 3, eGFP HEK293T media was aspirated and replaced with 1 mL NK92 cell culture. On day 4 cells were analyzed via flow cytometry.

### XXI. Mass Spec Assay

#### Generation of IgG subtype specific peptides

To establish unique IgG peptides, a tryptic digest of purified IgG subclasses (purchased from Rockwell) was prepared. 5 μg of each protein were digested using ProtiFi S-Trap micro columns (see details below). The resulting peptides were dried in a vacuum centrifuge. Prior to their analysis, they were resuspended in 50 μL 3% ACN, 0.1% FA and diluted 1:10. The peptides were separated on a nanoElute2 UHPLC system (Bruker Daltonics) and analyzed with diaPASEF on a timsTOF HT (Bruker Daltonics) (see details below). The measurements were searched using Spectronaut v. 20.1 (Biognosys) in directDIA mode using the default settings and cross-run normalization disabled.

The digests were compared and subtype-specific peptides were defined based on the comparisons and previous literature ^70, 71^. The selected peptides were VNSAAFPAPIEK (IgG1), DLPSPIER (IgG2b), ALPSPIEK (IgG2c), and ALPAPIER (IgG3). As these peptides could be confidently and reproducibly identified, heavy versions (AQUA Ultimate) with labeled lysine or arginine residues were custom synthesized and ordered from Thermo Fisher Scientific.

#### IgG pulldown

For each mouse sample, 100 μL of protein G magnetic beads (Thermo Fisher) were added to a 100 μL of binding/wash buffer (Thermo Fisher) in a 1.5 mL microcentrifuge tube (Protein Lobind, Eppendorf). Using a magnetic rack to separate the beads from the buffer, the buffer was exchanged with 1 mL of binding/wash buffer and washed for 1 min before buffer removal. A sample mixture (containing 15 μL of murine serum and 485 μL binding/wash buffer) was then added to the washed beads and incubated 1 hr (on a rotating shaking set at 1000 rpm and 22 ° C). The supernatant was then removed, and the beads were washed 3x with high ionic strength washes (1x TBS + 0,05% Tween-20) and 2x low ionic washes (20 mM Tris +150mM NaCl + 0.1 % Tween-20). The IgG’s were then eluted with 100 μL of elution buffer (Thermo Fisher, 21004) for 10 min with occasional mixing. After discarding the beads, 15 μL of neutralization buffer (1M Tris, pH 8.5) were added to IgG eluents.

100 μL of IgG eluate was prepared for mass spectrometry using the S-Trap Micro Universal Proteomics Sample prep kit from Protifi (Protifi, K02-micro-160) according to the manufacturer’s instructions. Samples were washed three times with solution 5 (step 11 in protocol) prior to digestion. For the digestion step, samples were incubated with 125 μL Trypsin/Lys-C mixture (Promega, V5073) diluted in solution 6. During digestion samples were first incubated at 37 °C for 30 min, and then at 47 °C for 1.5 hrs. Samples were eluted according to manufacturer’s protocol and dried in a vacuum centrifuge. One test sample was resuspended in 50 μL 3% ACN, 0.1% FA, diluted 1:20 and measured to estimate the endogenous levels of IgG peptides. The levels were determined to be at ∼ 2.5-5 fmol/ μL and therefore, all samples were resuspended in 50 μL 3% ACN, 0.1% FA and diluted 1:10 in a mixture of a tryptic digest of bovine serum albumin (BSA; Bruker 8217498; 50 fmol/ μL), iRT peptides (Biognosys/Bruker, 1816351), and a mixture of heavy peptides at a final concentration of 5 fmol/uL. For the calibration curves, heavy peptide dilutions with 50 fmol/uL, 25 fmol/uL, 12.5 fmol/μL, 5 fmol/μL, 2.5 fmol/μL, 500 amol/uL, 50 amol/uL, 5 amol/uL, and 500 zmol/μL were prepared (in the same BSA + iRT peptide background).

#### LC-MS/MS

1 μL of each sample was injected onto a nanoElute2 UHPLC system (Bruker Daltonics). The sample was loaded onto a trap cartridge (5 mm PepMapTM Neo Trap Cartridge, Thermo Scientific), washed, and then separated on a PepSep XTREME C18 25cmx150um, 1.5 μm column (Bruker Daltonics) using a 25-minute method. Peptides were separated with a 20 minute gradient from 0-26% B (eluent A = 3% ACN, 0.1% FA in water; eluent B = 0.1% FA in ACN) over 15 min, then from 26-32% B over an additional 5 min, followed by a 95% B wash, at a flow rate of 1 uL/min The eluted peptides were measured on a timsTOF HT mass spectrometer (Bruker Daltonics) using a diaPASEF method with a mobility window ranging from 0.7-1.3 1/K0, a ramp/accumulation time of 75 ms, and 38 equally sized 25 Da MS/MS windows, resulting in a cycle time of 0.81 s.

#### Data analysis

All measurements were analyzed in Skyline and R using the R packages tidyverse^90^ and data.table^91^. iRT peptides were used to check for retention time inconsistencies. To produce calibration curves, the calculated areas of the 3-5 best ranked fragments were summed up for each sample. After assessing the linearity and excluding lower data points that were not within the linear range, a linear model (lm) was fit onto the log10 transformed, summed up area values at the respective log10 transformed concentrations. Slope and intercept were calculated for each heavy labeled peptide. The limit of detection (LOD) and the limit of quantification (LOQ) were calculated from 3 blank samples and 3 BSA peptide + iRT injections as follows:

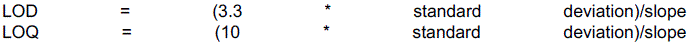

The light peptide abundances were normalized in regards to the heavy spiked in peptides (at 5 fmol/uL) to account for any sample losses or inconsistencies. The normalized light intensities were calculated as follows: Correction factor = expected heavy (calculated from calibration curve) / measured heavy Normalized light = intensity light * correction factor

The previously calculated slope and intercept values were used to calculate the concentrations. Relative abundance values were calculated using the determined calculations. Specifically, abundances were calculated by normalizing all IgG-subtype-specific peptide levels to the mean concentration of peptides from the AAV IgG κ nanobody condition.

The mass spectrometry proteomics data, as well as all relevant search and analysis files, have been deposited to the ProteomeXchange Consortium via the PRIDE^92^ partner repository with the dataset identifier PXD079901.

### XXII. Collagen disease model

Clearance assay: C57BL/6 mice were injected via tail vein with 1e11 viral genomes of AAV expressing IgG κ mGELYTAC or controls. A weeks later, 100 mg/kg fluorescently labeled α collagen antibody cocktail (MDbioproducts CIA-MAB-2C), and serum was collected and analyzed as described above.

Inflammation model: Methods were based on previously published protocols^72^. In short, C57BL/6 mice were first injected with 200 mg/kg α collagen antibody cocktail intraperitoneally and then an hour later injected with 10 mg/ kg efgartigimod alfa or 1e11 viral genomes of AAV (encoding IgG κ mGELYTAC or IgG κ nanobody) via tail vain. 2 days later, 100μg of lipopolysaccharide (200 μL of 0.5 mg/mL) was injected intraperitoneally to induce further inflammation.

### XXIII. Disease model scoring

Mouse paws were scored blindly by two blinded reviewers according to the following rubric: each swollen toe/ finger was given a score 1. Each toe/finger that is beginning to swell or has receding swelling was given a score of 0.5. For each palm or ankles, a score of 1 (beginning to swell/ receding swelling), 2 (fully swollen), or 3 (ankylosed). Mice were excluded from the study if they did not hit a minimum disease score of 15. Paw images were taken using a Canon Rebel T2i (2010) with an EF 100mm f/2.8L Macro IS USM macro lens.

### XXIV. Rotarod assay

Mice were first conditioned for 3 days prior to rotarod measurements. For rotarod measurements (LabeoTech, 38305), mice were placed onto the rotarod apparatus. The rotarod settings were as follows: gradual ramp from 0-30 RPM for 180 seconds. An auditory clap was given before the start of the rotarod experiment.

## Acknowledgments

We thank Prof. Albert Heck, Mirat Sojitra, Sophia Shi, Veronica Li, Johannes Hevler, and Nick Till for protocols and helpful discussions. We thank Prof. Simran Johal, Dr. John S. Tamaresis, and School of Medicine Data Studio for help with statistical analysis. The Data Studio is supported by the National Center for Advancing Translational Sciences of the National Institutes of Health under Award Number UM1TR004921. The content is solely the responsibility of the authors and does not necessarily represent the official views of the National Institutes of Health. C.L.M., S.A.Y.-H. and L.L. are members of the Parker Institute for Cancer Immunotherapy, which supports the Stanford University Cancer Immunotherapy Program. C.L.M. acknowledges support from the Virginia and D.K. Ludwig Fund for Cancer Research. We also thank Chandan Kadur and Ki Eun Pyo for cell culture assistance. Fig. 2g was made using templates from https://BioRender.com. Mouse figures are made from NIAID NIH BIOART (source: bioart.niaid.nih.gov/bioart/279).

## Funding

G. Harold & Leila Y. Mather’s Charitable Foundation grant MF-2503-09193 (C.R.B)

National Institutes of Health grant R01GM058867 (C.R.B.)

NexTGen team supported by the Cancer Grand Challenges partnership funded by:

Cancer Research UK CGATF-2021/100005 (C.R.B)

National Cancer Institute NIH/OT2CA278686 (C.R.B)

The Mark Foundation for Cancer Research (C.R.B)

National Science Foundation grant 2037164 (K.D., and C.R.B.)

BioX SIGF Fellowship (J.L.Y.)

BioX Bowes Fellowship (K.Y.L.)

Sarafan ChEM-H Chemistry/Biology Interface Predoctoral Training Program (K.Y.L)

Sarafan ChEM-H Chemistry/Biology Interface Predoctoral Training Program (C.S.E.)

Parker Institute for Cancer Immunotherapy Parker Scholarship (S.A.Y.-H.)

## Author contributions

J.L.Y., K.Y.L., and C.R.B. conceived the idea of the study. J.L.Y., K.Y.L., C.S.E., D.S., S.A.Y.-H., and L.L. planned experiments and conducted the investigation. K.D., S.A.Y.-H., L.L., E.S., and C.L.M. contributed new reagents/analytic tools. K.D. and C.R.B. acquired funding for the project and supervised J.L.Y., K.Y.L., and C.S.E. The initial manuscript was written by J.L.Y. and C.R.B. with input and reviewing from K.Y.L., C.S.E., S.A.Y.-H., L.L., E.S., C.L.M., and K.D.

## Corresponding Author

**Extended Data Fig. 1.**
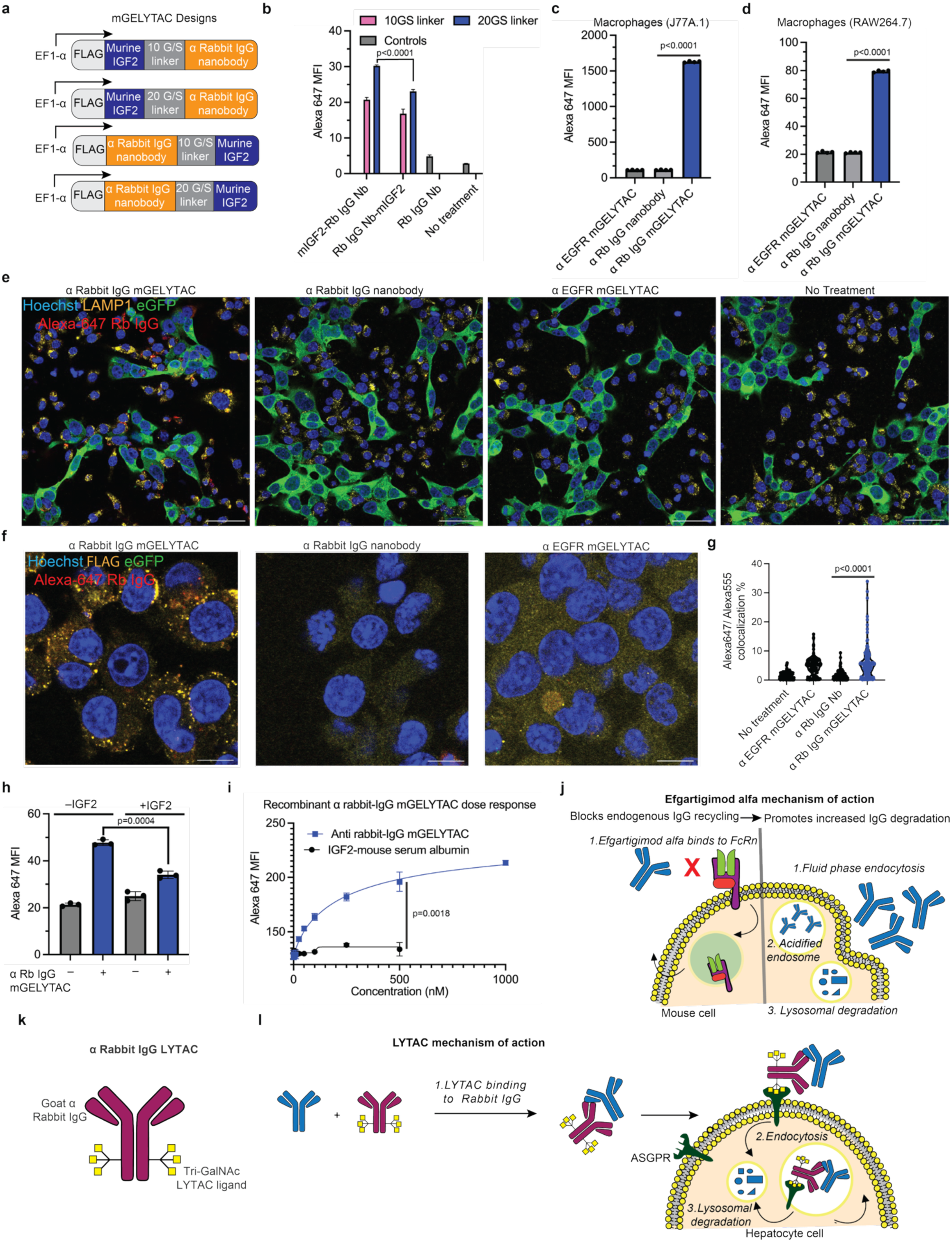
mGELYTAC mediate uptake IgG in murine cell culture & *in vivo.* (a) Schematic of mGELYTACs constructs utilizing 2 different orientations (N or C terminal mIGF2 fusion) and 2 different linker lengths (10 or 20 glycine/serine linkers). (b) Alexa-647 median fluorescence from X16C8.5 mouse lymphocytes cocultured with GFP HEK293T cells secreting α rabbit IgG mGELYTACs (4 different mGELYTACs from Extended Data Fig. 1a) or rabbit-IgG nanobody control. Cocultured cells were treated with Alexa-647 rabbit-IgG (100 nM) for 24 hours. Then, the cell mixture was lifted and analyzed using flow cytometry. Error bars represent the SD from 3 biological replicates. p values were determined using Welch’s t-test. (c and d) Alexa-647 median fluorescence from J77A macrophages (c) or RAW 264.7 (d) cocultured with eGFP HEK293T cells secreting α rabbit IgG mGELYTAC, rabbit-IgG nanobody control, or a control mGELYTAC (targeting human EGFR). Cocultured cells were treated with Alexa-647 rabbit-IgG (100 nM) for 24 hours. Then, the cell mixture was lifted and analyzed using flow cytometry. Error bars represent the SD from 4 biological replicates. p values were determined using Welch’s t-test. (e) Expanded field view of images shown in Fig. 1j. Scale bars, 50 μm. (f) Confocal fluorescence images of cocultured eGFP HEK23T cells secreting α rabbit IgG mGELYTAC, rabbit-IgG nanobody control, or a control mGELYTAC (targeting human EGFR) were cocultured with murine J77A macrophages. Cocultured cells were treated with Alexa-647 rabbit-IgG (100 nM) for 24 hours on coverslips. Then, the cell mixture was fixed and stained with an antibody against FLAG (mGELYTAC) and Hoechst (nuclear stain). Scale bars, 10 μm. (g) Quantification of confocal images to show colocalization of Alexa-647 rabbit-IgG with FLAG (mGELYTAC) staining. The quantification is defined as pixels within the image that contain both Alexa-647 (rabbit IgG) and Alexa-555 (FLAG) signal. Error bars represent the SD from 114-150 cells. p values were determined using Mann-Whitney U test. (h) Alexa-647 MFI (median fluorescence intensity) of MC38 mouse epithelial cells cocultured with eGFP HEK293T cells secreting α rabbit IgG mGELYTAC. Cocultured cells were treated recombinant mouse IGF2 (3.4 μM) and Alexa-647 rabbit-IgG (100 nM) for 24 hours. Then, the cell mixture was lifted and analyzed using flow cytometry. Error bars represent the SD from 3 biological replicates. p values were determined using Welch’s t-test. (i) Alexa-647 rabbit IgG internalization as a function of mGELYTAC concentration. MFI (median fluorescence intensity) of X16C8.5 cells was measured 24 hr after treatment with recombinant α rabbit IgG mGELYTAC (squares) and AF647-rabbit IgG (100 nM). Negative control is IGF2-MSA (circles). Errors bars represent the SD from three biological replicates. p value determined using Welch’s t-test. (j) Schematic depicting efgartigimod alfa mechanism of action. (k) Schematic showing α Rb LYTAC design^22^. (l) Schematic outlining α Rb LYTAC mechanism of action.

**Extended Data Fig. 2.**
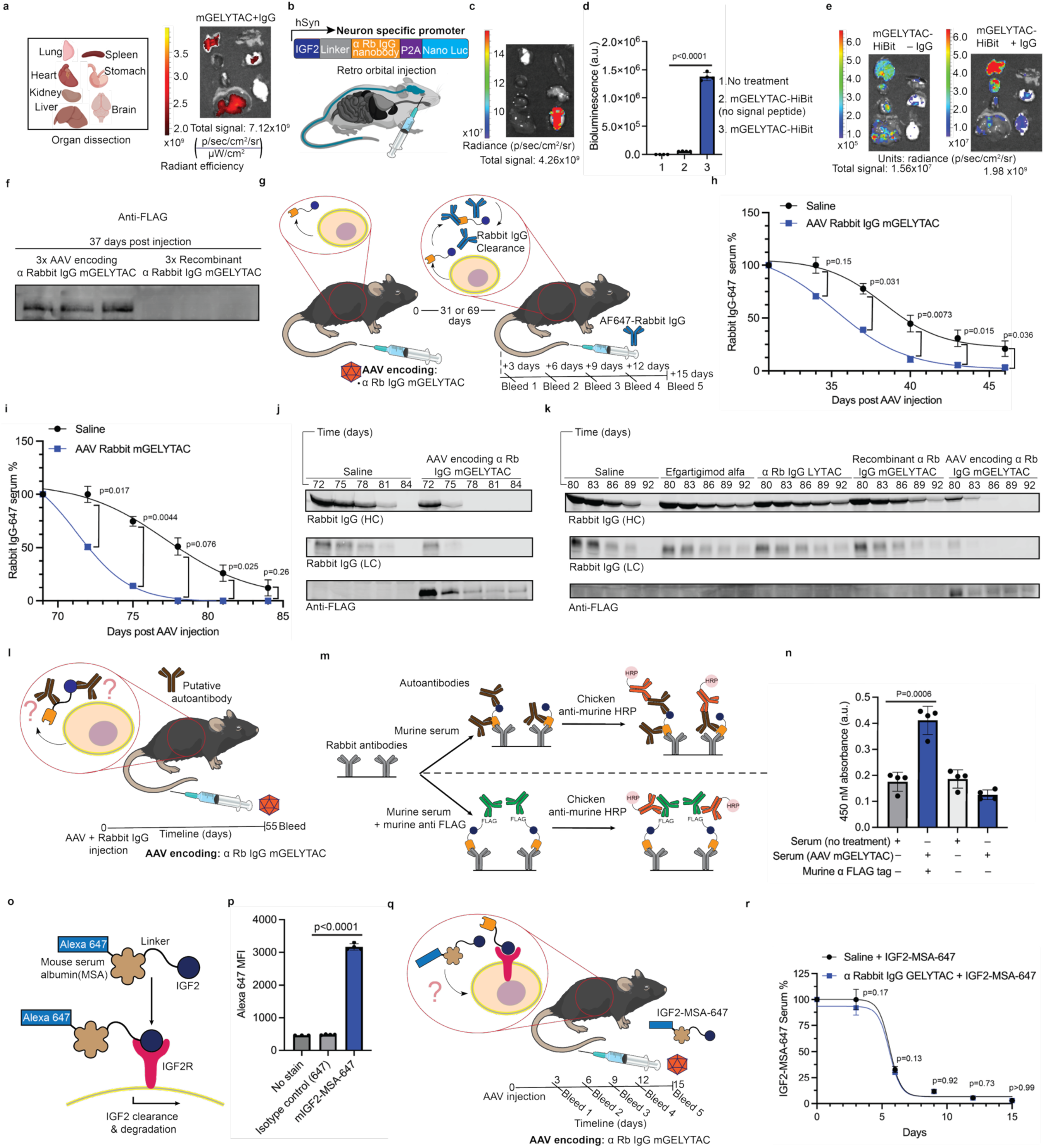
Gene therapy delivered mGELYTACs clear exogenous IgGs *in vivo.* (a) Alexa-647 signal from mouse organs from a mouse treated with AAV8 encoding mGELYTAC and Alexa-647 rabbit IgG (40 mg/kg) for 6 days. The organ arrangement schematic represents the order organs in all organ imaging in this figure. (b) Schematic for mGELYTAC under the control of the neuron specific promoter hSyn. This construct includes nano luciferase downstream of the mGELYTAC to enable bioluminescence and is packaged using AAV.PhP.eb serotype. (c) Bioluminescence from organs from mice treated with AAV.PhP.eb encoding mGELYTAC-P2A-nLuc at 3 days. (d) Bioluminescence from untreated HEK293T cells or HEK293T cells transfected with mGELYTAC-HiBit (no signal peptide control) or mGELYTAC-HiBit. Error bars represent the SD from 4 biological replicates. p values were determined using Welch’s t-test. (e) A comparison of bioluminescence from organs from mice treated with AAV8 encoding mGELYTAC-HiBit at day 6 with or without 40 mg/kg rabbit IgG. (f) α FLAG western blot of murine plasma from mice treated either recombinant α Rb IgG mGELYTAC or AAV8 encoding for α Rb IgG mGELYTAC. (g) Schematic outlining experiment to monitor mGELYTAC longevity. C57BL/6 mice were injected with AAV viral particles encoding for rabbit IgG targeting mGELYTACs. 31 or 69 days later, mice were injected with 15 mg/kg Alexa-647 labeled rabbit IgG. Submandibular bleed was performed every 3 days from day 3-15 post IgG administration. Analysis was performed with plate reader (31 day and 69 day) and western blot (69 day). (h) Alexa-647 rabbit IgG serum clearance as a function of time for mice treated with AAVs encoding for α rabbit IgG mGELYTAC (squares), 31 days prior to AF647-rabbit IgG administration. Rabbit IgG clearance is measured via Alexa-647 plate reader assay on samples collected between 34-46 days. Error bars represent the SEM from 4 biological replicates. p values were determined using Welch’s t-test. (i) Alexa-647 rabbit IgG serum clearance as a function of time for mice treated with AAVs encoding for α rabbit IgG mGELYTAC (squares), 69 days prior to AF647-rabbit IgG administration. Rabbit IgG clearance is measured via Alexa-647 plate reader assay on samples collected between 72-84 days. Error bars represent the SEM from 3 biological replicates. p values were determined using Welch’s t-test. (j) Western blot analysis of Alexa-647 rabbit IgG and mGELYTAC levels in longevity experiment outlined in Extended Data Fig. 2g,i. Alexa-647 rabbit IgG levels were determined via IR700 measurement. Labeled heavy chains appear at ∼50 kDa and labeled light chains appear at ∼25 kDa. α FLAG staining was also performed to analyze mGELYTAC levels. (k) Western blot analysis of Alexa-647 rabbit IgG and mGELYTAC levels in AAV delivered mGELYTAC and IgG depleting biologics comparison outlined in Fig. 2k. Alexa-647 rabbit IgG levels were determined via IR700 measurement. Labeled heavy chains appear at ∼50 kDa and labeled light chains appear at ∼25 kDa. α FLAG staining was also performed to analyze mGELYTAC levels. (l) Schematic outlining assay to detect putative autoantibodies. AAV viral particles encoding for rabbit IgG targeting mGELYTACs were injected via tail vein. A submandibular bleed was performed at day 55, and an ELISA was performed to detect autoantibodies. (m) Schematic outlining ELISA to detect putative autoantibodies against mGELYTACs targeting rabbit IgG. Plates were coated with rabbit antibodies and incubated either with murine plasma (no treatment or AAV encoding rabbit IgG targeting mGELYTAC) or murine plasma (no treatment or AAV encoding rabbit IgG targeting mGELYTAC) with murine α FLAG antibody. The plasma co-incubated with murine α FLAG is a mock autoantibody, since mGELYTACs contain a FLAG tag. After initial incubation and a series of washes, the plate is treated with chicken α mouse HRP conjugate and subsequently incubated with HRP substrate and readout on plate reader. (n) 450 nM absorbance data from ELISA outlined in (m). Error bars represent the SD from 4 biological replicates. p value determined using Welch’s t-test. (o) schematic depicting IGF2-MSA (mouse serum albumin)-Alexa-647 binding to IGF2R. (p) Alexa-647 median fluorescence intensity from unstained MC38 cells or MC38 cells stained with Alexa-647 IGF2-MSA (mouse serum albumin) or mouse Alexa-647 IgG isotype control. Error bars represent the SD from three to four biological replicates. p value determined using Welch’s t-test. (q) Schematic outlining experiment to assess IGF2 clearance in mice treated with AAV8 encoding α rabbit IgG mGELYTAC or saline. IGF2-MSA-AF647 (5.8 mg/kg) was co-injected with viral treatment. Submandibular bleeds were performed every 3 days from days 3-15 of AAV/IGF2-MSA-647 administration and before analysis with plate reader (r) IGF2-MSA-647 serum clearance as a function of time for mice treated with AAVs encoding for α rabbit IgG mGELYTAC (squares) or saline (circles). IGF2-MSA-647 is measured via Alexa-647 plate reader assay on samples collected between 3-15 days. Error bars represent the SEM from 4 biological replicates. p values were determined using Welch’s t-test.

**Extended Data Fig. 3:**
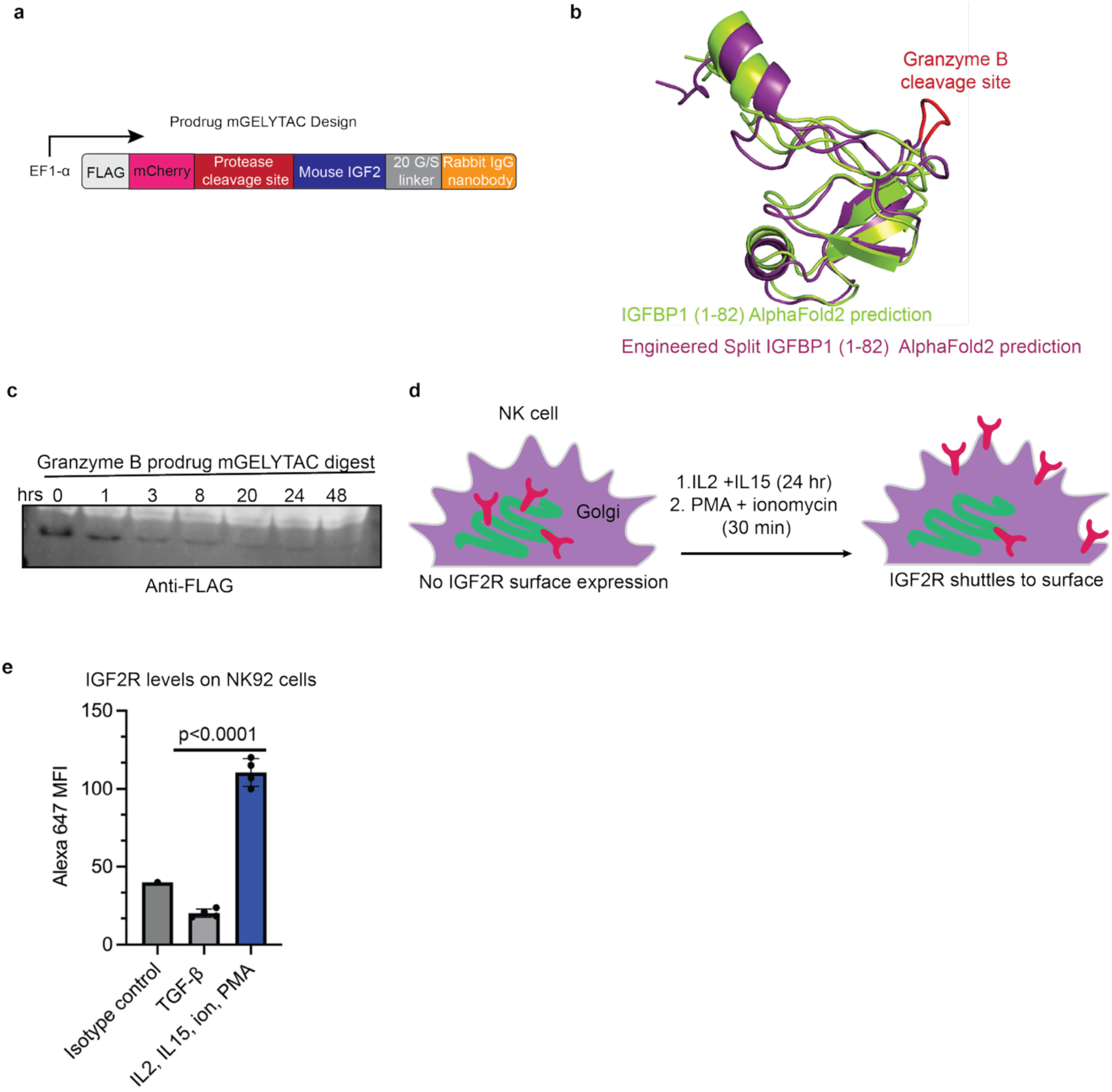
Development of protease-activatable mGELYTACs. (a) Schematic of prodrug mGELYTAC control, where the IGFBP group caging IGF2 is replaced with mCherry. mCherry has no known affinity to IGF2 and provides only steric hindrance to IGF2. (b) Predicted split IGFBP structure containing granzyme B (GzmB) cleavage site (red). Predicted structure was generated using AlphaFold2. (c) α FLAG western blot showing GzmB gated mGELYTAC levels incubated with recombinant GzmB (375 nM) over the course of 48 hrs. (d) Schematic outlining hypothesized IGF2R translocation upon treatment with immune-activating conditions. (e) Alexa-647 MFI (median fluorescence intensity) representing IGF2R levels from NK92 cells either treated with TGF-β (immunosuppressive) or IL2 + IL15 + ionomycin + PMA (immune-activating). Post treatment, cells were stained with an α IGF2R-647 antibody and analyzed via flow cytometry. Error bars represent the SD from four biological replicates. p value determined using Welch’s t-test.

**Extended Data Fig. 4:**
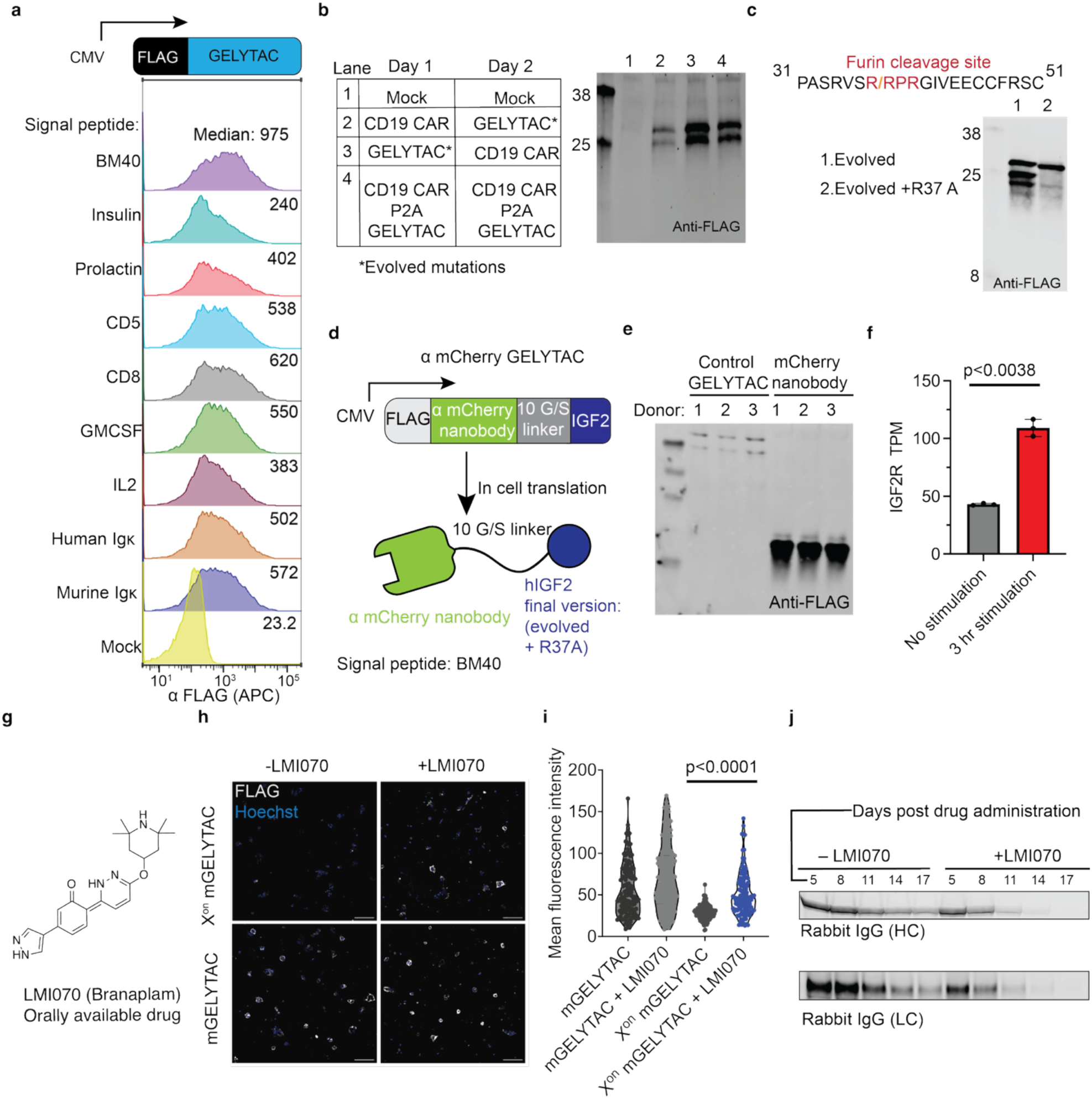
Cell-based synthetic biology applications of GELYTACs. (a) Flow cytometry of primary T-cells infected with virus encoding GELYTACs downstream of various signal peptides, stained with α FLAG antibody. Signal peptides were selected from proteins with high expression in T-cells. Although not tethered to cell, GELYTACs have high back-binding and can be stained. (b) α FLAG western blot of T-cell supernatant infected with differing orders of GELYTAC (evolved mutations^31^) and CD19 CAR retroviruses. T-cells were either infected on day 1 with GELYTAC and day 2 with CAR, day 1 with CAR and day 2 with GELYTAC, or day 1 and 2 with virus encoding for CAR followed by a ribosomal skipping P2A sequence and then GELYTAC. (c) Protease sequence analysis around the 31 and 51 region reveals a furin cleavage site^63^, and the corresponding mutation R37A^63^ rescues GELYTAC from cleavage in α FLAG western blot of supernatant transfected HEK293T cells. (d) Schematic outlining optimized α mCherry GELYTACs for CAR-T applications. (e) α FLAG western blot from CAR-T cell supernatant of controls from 3 donors used in coculture experiment depicted in Fig. 4a. (f) IGF2R transcriptomics from CD19 CAR-T cells either unstimulated or stimulated for 3 hrs. This data set is previously published and is publicly availbile^65^. Error bars represent the SD from three biological replicates (three donors). p value determined using Welch’s t-test. (g) Chemical structure of LMI1070 (Branaplam). The orally available small molecule used in drug induced splicing experiments. (h) Confocal fluorescence images of HEK23T cells secreting α rabbit IgG mGELYTAC under X^on^ control (+/− LMI070), and regular α rabbit IgG mGELYTAC without X^on^ (+/− LMI070). Cells were plated and treated on coverslips. The cell mixture was then fixed and stained with an antibody against FLAG (mGELYTAC) and Hoechst (nuclear stain). Scale bars, 100 μm. (i) Quantification of mean fluorescence intensity corresponding FLAG staining in confocal images in Extended Data Fig. 4h. The quantification is defined as pixels within the image that contain both Hoechst staining and FLAG-tag staining (FLAG). Error bars represent the SD from 103-140 cells. p value determined using Mann-Whitney U non-parametric test. (j) Western blot analysis of Alexa-647 rabbit IgG levels expression in drug induced splicing experiment outlined in Fig. 4g**-i** Alexa-647 rabbit IgG levels were determined via IR700 measurement. Labeled heavy chains appear at ∼50 kDa and labeled light chains appear at ∼25 kDa.

**Extended Data Fig. 5.**
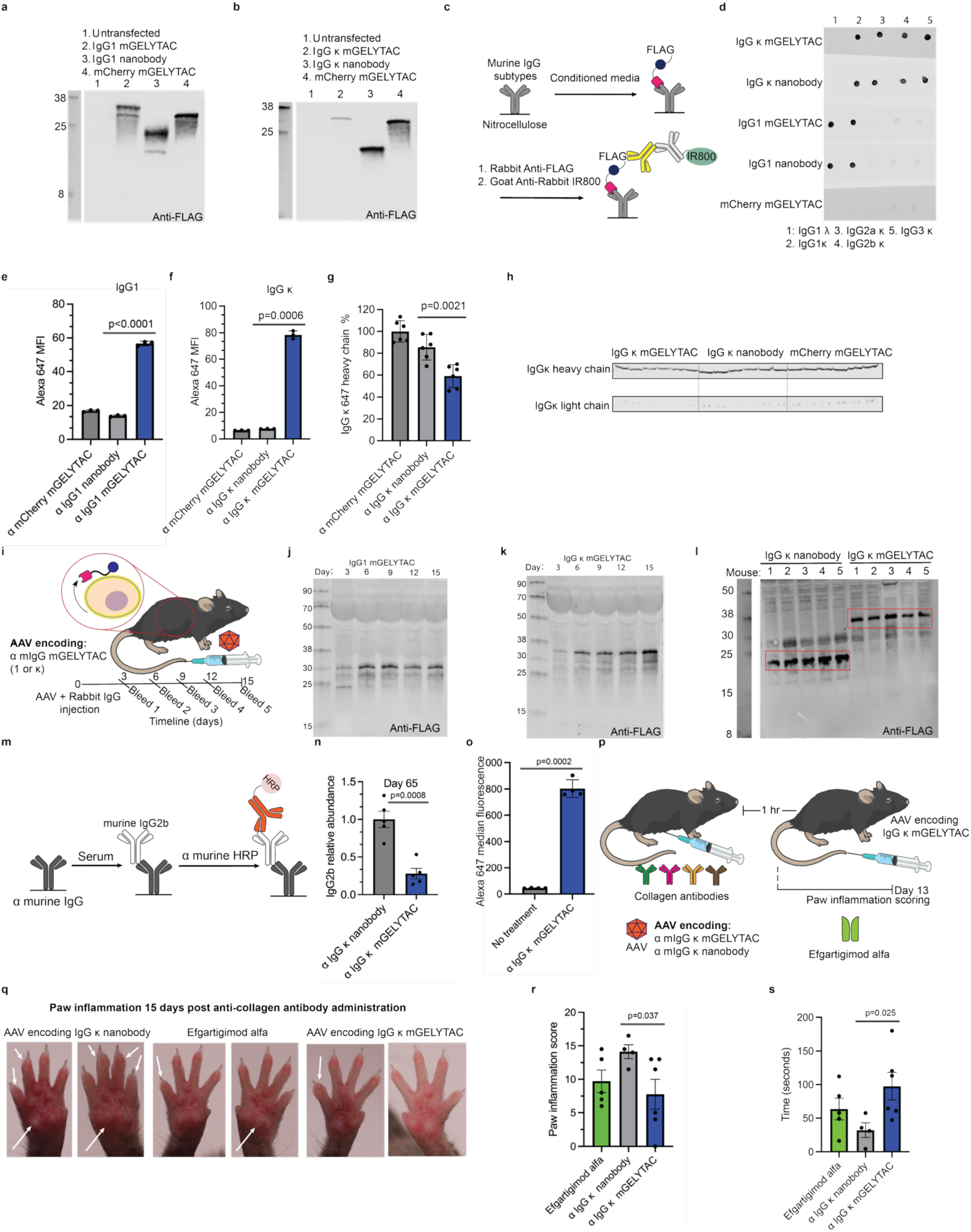
Gene therapy delivered mGELYTACs clear endogenous murine IgG. (a) α FLAG western blot of conditioned supernatant from eGFP HEK293T cells transiently transfected with α murine IgG1 mGELYTACs, α murine IgG1 nanobody only, and a non-targeting mGELYTAC control (targeting mCherry^69^). (b) α FLAG western blot of conditioned supernatant from eGFP HEK293T cells transiently transfected with α murine IgG κ mGELYTACs, α murine IgG κ nanobody only, and a non-targeting mGELYTAC control (targeting mCherry). (c) Schematic of a dot blot analyzing murine IgG targeting mGELYTAC selectivity. (d) Dot blot described in (c). (e and f) Median Alexa-647 fluorescence of X16C8.5 cells cocultured with eGFP HEK293T cells secreting α murine IgG mGELYTAC (IgG1 or IgG κ), IgG nanobody control (IgG1 or IgG κ), or a control mGELYTAC (targeting mCherry). Cocultured cells were treated with Alexa-647 mouse-IgG1 (100 nM) for 24 hours. Then, the cell mixture was analyzed using flow cytometry. Error bars represent the SD from four biological replicates. p value determined using Welch’s t-test. (g) To quantify degradation of IgG κ, after addition of 50 nM Alexa 647-labeled IgG κ for 48 hours, both cells and media were collected and analyzed for Alexa-647 signal via blot. This experiment was repeated six times. The control is HEK293T cells transfected either IgG κ nanobody or mCherry mGELYTAC. Error bars represent the SD from six biological replicates. p value determined using Welch’s t-test. (h) Western blot analysis of Alexa-647 murine IgG κ levels in eGFP HEK293T and X16C8.5 coculture depicted in Extended Data Fig. 5g. Alexa-647 murine IgG levels were determined via IR700 measurement. Labeled heavy chains appear at ∼50 kDa and labeled light chains appear at ∼25 kDa. (i) Schematic outlining AAV infection experiment. AAV viral particles encoding for murine IgG targeting mGELYTACs were injected via tail vein. Submandibular bleeds were performed every 3 days from days 3-15 and then analyzed via α FLAG western blot. (j) α FLAG western blot of murine plasma collected from C57BL/6 mice infected with AAV8 encoding for IgG1 mGELYTAC. Murine plasma was collected every 3 days from days 3-15. (k) α FLAG western blot of murine plasma collected from C57BL/6 mice infected with AAV8 encoding for IgG κ mGELYTAC. Murine plasma was collected every 3 days from days 3-15. (l) α FLAG western blot of murine plasma collected from C57BL/6 mice infected with AAV8 encoding for IgG κ nanobody or mGELYTAC 65 days prior. 5 mice for each condition shown. (m) Schematic depicting ELISA to monitor IgG2b depletion. (n) IgG2b plasma relative abundance from C57BL/6 mice treated 65 days prior with AAV8 encoding IgG κ mGELYTACs or nanobody determined from ELISA. Error bars represent the SEM from 5 biological replicates. p value determined using Welch’s t-test. (o) Median Alexa-647 fluorescence of X16C8.5 cells cocultured with eGFP HEK293T cells secreting α murine IgG κ mGELYTAC. Cocultured cells were treated with Alexa-647 labeled α collagen antibody cocktail (100 nM) for 24 hours. Then, the cell mixture was analyzed using flow cytometry. Error bars represent the SD from four biological replicates. p value determined using Welch’s t-test. (p) Schematic outlining arthritis experiment. α Collagen antibodies were first passively transferred to mice (200mg/kg). Then 1 hr later, mice were injected with AAV encoding for α IgG κ mGELYTAC, AAV encoding α IgG κ nanobody, or efgartigimod alfa. Mice were then scored 13 days post treatment. (q) Images of mouse paws given the treatment outlined in (p). (r) Paw inflammation scores from mice given the treatment outlined in (p). Scores are averages from two blinded reviewers. Error bars represent the SEM from 4 (nanobody), 5 (efgartigimod alfa), or 6 (mGELYTAC) biological replicates. p values were determined using Welch’s t-test. (s) Time mice remained on spinning rotarod apparatus 8 days post mGELYTAC AAV treatment. Error bars represent the SEM from 4 (nanobody), 5 (efgartigimod alfa, or 6 (mGELYTAC) biological replicates. p values were determined using Welch’s t-test.

## Supplementary information

Contents:

1. Flow cytometry gating
2. Recombinant biologics production
3. mCherry cell line validation

## Flow cytometry gating

**Supplementary Figure 1.**
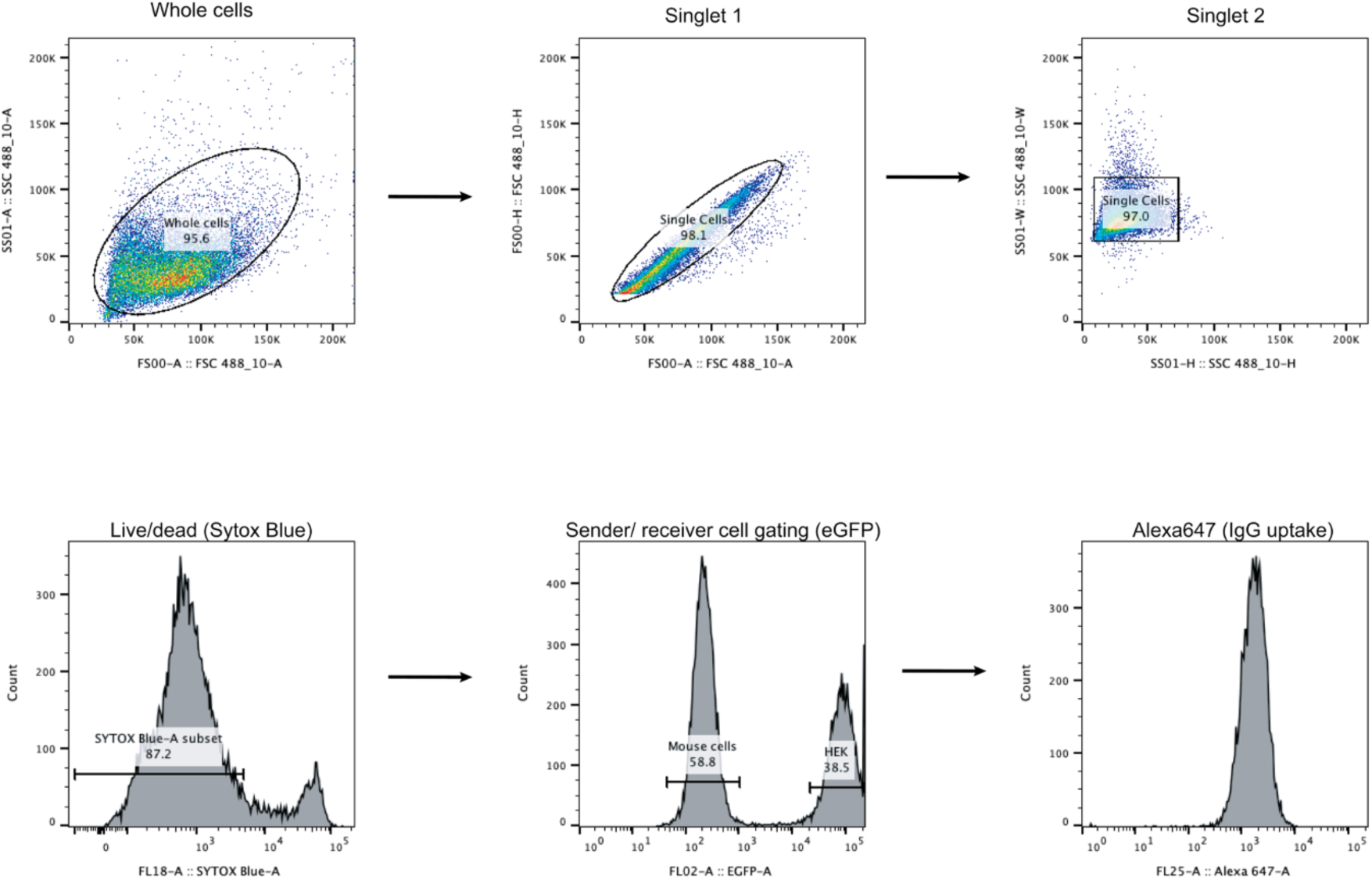
Flow cytometry gating scheme.

## Recombinant biologics production

**Supplementary Figure 2.**
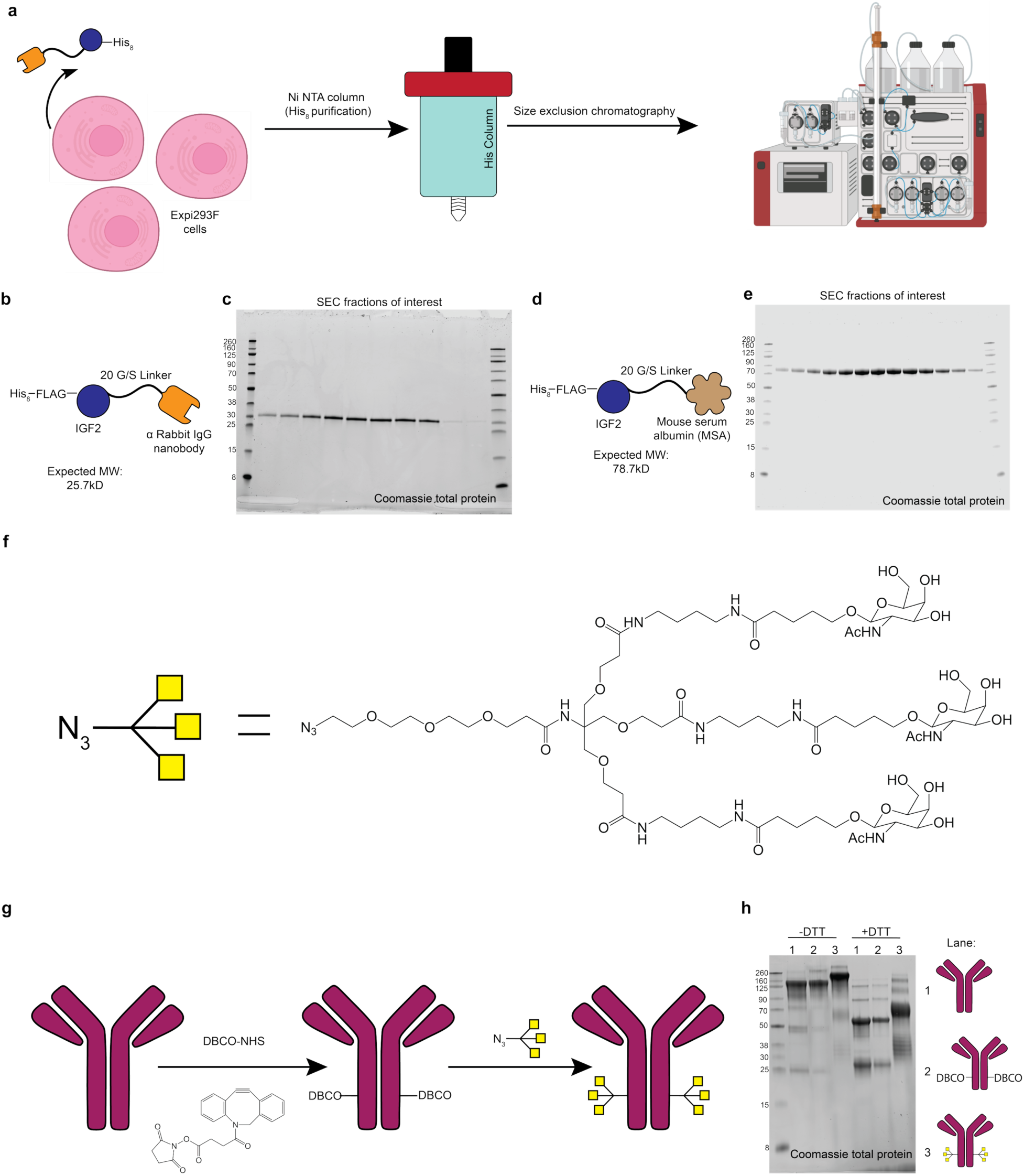
(a) Schematic showing process where recombinant mGELYTAC or IGF2-MSA proteins were made. (b) Schematic of recombinant α rabbit IgG mGELYTAC design. (c) Coomassie stained gel of size exclusion chromatography (SEC) fractions for α rabbit IgG mGELYTAC purification. (d) Schematic of recombinant IGF2-MSA design. (e) Coomassie stained gel of SEC fractions from IGF2-MSA purification. (f) Schematic of GalNAc LYTAC ligand. (g) Schematic for LYTAC bioconjugation. (h) Coomassie stained gel (with or without reducing conditions) of unconjugated goat α rabbit IgG, DBCO labeled goat α rabbit IgG, or complete LYTAC-antibody conjugate.

**Supplementary Figure 3.**
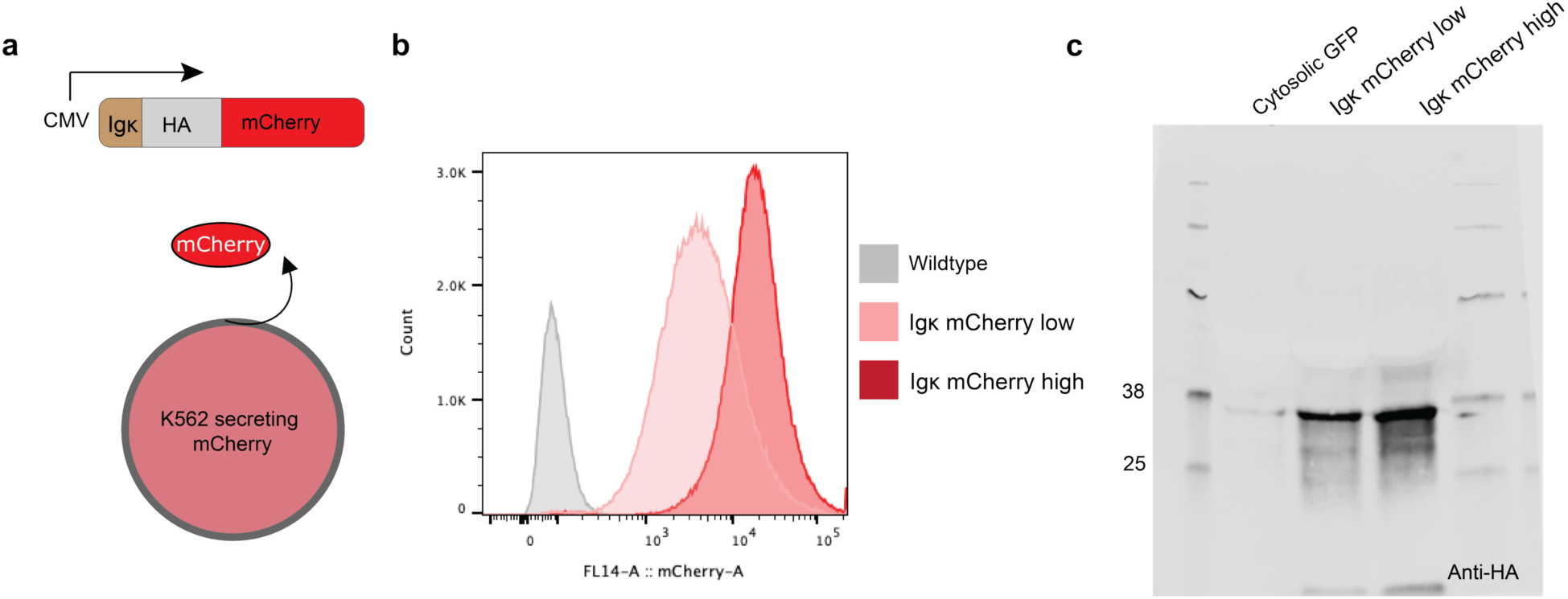
(a) Schematic of mCherry secreting K562 cell line used in assays in Fig. 4 a,c,d,e. (b) Flow cytometry analysis of mCherry secreting cell lines. Igk mCherry high cell line was used in the final assay. (c) Anit-HA western blot of supernatant collected from mCherry secreting K562s or control (expressing cytosolic GFP).

## References

1. Nimmerjahn, F., Vidarsson, G. & Cragg, M. S. Effect of posttranslational modifications and subclass on IgG activity: from immunity to immunotherapy. Nat. Immunol. 24, 1244–1255 (2023).

2. Hoi, A., Igel, T., Mok, C. C. & Arnaud, L. Systemic lupus erythematosus. Lancet 403, 10441, 2326–2338 (2021).

3. Kelley, B. P. et al. Autoimmune encephalitis: Pathophysiology and imaging review of an overlooked diagnosis. *Am*. J. Neuroradiol. 38, 6, 1070–1078 (2017).

4. Smolen, J. S., et al. Rheumatoid arthritis. Nat. Rev. Dis. Primers 4,18001 (2018).

5. Di Matteo, A., Bathon, J. M. & Emery, P. Rheumatoid arthritis. Lancet 402, 10416, 2019–2033 (2023).

6. Zorn, E. & See, S. B. Is there a role for natural antibodies in rejection following transplantation? Transplantation 103,8, 1612–1619 (2019).

7. Schirmer, D. A. et al. Mouse models of IgG-and IgM-mediated hemolysis. Blood 109, 7,3099–3107 (2007).

8. Ma, S., et al. Spatial transcriptomic landscape unveils immunoglobin-associated senescence as a hallmark of aging. Cell 187, 24, 7025–7044 (2024).

9. Stockfelt, M., Teng, Y. K. O. & Vital, E. M. Opportunities and limitations of B cell depletion approaches in SLE. Nat Rev Rheumatol. 21,111–126 (2025).

10. Zanatta, E., Cozzi, M., Marson, P. & Cozzi, F. The role of plasma exchange in the management of autoimmune disorders. Br. J. Haematol. 186, 2, 207–219 (2019).

11. Wyckoff, S. L. & Hudson, K. E. Targeting the neonatal Fc receptor (FcRn) to treat autoimmune diseases and maternal-fetal immune cytopenias. Transfusion. 61, 5, 1350–1354) (2021).

12. Cutolo, M. & Straub, R. H. Sex steroids and autoimmune rheumatic diseases: state of the art. Nat. Rev. Rheumatol. 16, 628–644, (2020).

13. Ronchetti, S. et al. A Glance at the Use of Glucocorticoids in Rare Inflammatory and Autoimmune Diseases: Still an Indispensable Pharmacological Tool? Front. Immunol. 11, 613435 (2021).

14. West, E. E., Woodruff, T., Fremeaux-Bacchi, V. & Kemper, C. Complement in human disease: approved and up-and-coming therapeutics. Lancet 403 (**10424**), 392–405 (2024).

15. Fauconnier, A., Melis, M., Berenbeck, M., Pio, B. & Croisier, T. Trends in the drug target landscape for autoimmune diseases. Nat Rev Drug Discov. 24, 415–416, (2024).

16. Tanaka, Y., Luo, Y., O’Shea, J. J. & Nakayamada, S. Janus kinase-targeting therapies in rheumatology: a mechanisms-based approach. Nat. Rev. Rheumatol. 18, 3, 133–145 (2022).

17. Youssef, J., Novosad, S. A. & Winthrop, K. L. Infection Risk and Safety of Corticosteroid Use. Rheum. Dis. Clin. North Am. 42, 1,157–176 (2016).

18. Peter, H. H. et al. Targeting FcRn for immunomodulation: Benefits, risks, and practical considerations. J. Allergy Clin. Immunol. 146, 3, 479–491 (2020).

19. Salerno, M. et al. Anabolic androgenic steroids and carcinogenicity focusing on Leydig cell: a literature review. Oncotarget 9, 27, 19415–19426 (2018).

20. Karagas, M. R. et al. Non-melanoma skin cancers and glucocorticoid therapy. Br. J. Cancer 85, 5, 683–686 (2001).

21. Ruiz, R. & Kirk, A. “Chapter 97 – Long-Term Toxicity of immunosuppressive Therapy” in Transplantation of the Liver (Third Edition). Elsevier Inc, Philadelphia, PA (2015).

22. Ahn, G., Banik, S. M. & Bertozzi, C. R. Degradation from the outside in: Targeting extracellular and membrane proteins for degradation through the endolysosomal pathway. Cell Chem. Biol. 28, 7, 1072–1080 (2021).

23. Wells, J. A. & Kumru, K. Extracellular targeted protein degradation: an emerging modality for drug discovery. Nat Rev. Drug Discov. 23, 126–140 (2024).

24. Banik, S. M. et al. Lysosome-targeting chimaeras for degradation of extracellular proteins. Nature 584, 7820, 291–297 (2020).

25. Ahn, G. et al. LYTACs that engage the asialoglycoprotein receptor for targeted protein degradation. Nat Chem Biol 17, 937–946 (2021).

26. Caianiello, D. F. et al. Bifunctional small molecules that mediate the degradation of extracellular proteins. Nat. Chem. Biol. 17(**9**), 947–953 (2021).

27. Zhou, Y., Teng, P., Montgomery, N. T., Li, X. & Tang, W. Development of Triantennary N – Acetylgalactosamine Conjugates as Degraders for Extracellular Proteins. ACS Cent. Sci. 7(**3**), 499–506 (2021).

28. Kougentakis, C. et al. Catalytic degradation of circulating targets with FcRn-mediated cycling LYTACs. Preprint at https://www.biorxiv.org/content/10.1101/2025.01.12.632472v1 (2025).

29. Mullard, A. Extracellular targeted protein degrader removes antibodies in first test in humans. Nat Rev. Drug Discov. 23, 483–486 (2024).

30. Biohaven Reports Positive Degrader Data Achieving > 80% Sustained Reductions in Total IgG with Potential First-in-Class BHV-1300, Biohaven press release (2025).

31. Yang, J. L. et al. Directed evolution of genetically encoded LYTACs for cell-mediated delivery. Proc. Natl. Acad. Sci. U.S.A. 121, 13 (2024).

32. Pleiner, T., Bates, M. & Görlich, D. A toolbox of anti-mouse and anti-rabbit IgG secondary nanobodies. J. Cell Biol. 217, 3, 1143–1154 (2018).

33. Han, Y., Li, M., Qiu, F., Zhang, M & Zhang, Y.-H. Cell-permeable organic fluorescent probes for live-cell long-term super-resolution imaging reveal lysosome-mitochondrion interactions. Nat Commun 8, 1307 (2017).

34. Tintelnot, J. et al., Nanobody Targeting of Epidermal Growth Factor Receptor (EGFR) Ectodomain Variants Overcomes Resistance to Therapeutic EGFR Antibodies. Mol. Cancer Ther. 18(**4**), 823–833 (2019).

35. Heo, YA. Efgartigimod: First Approval. Drugs 82, 341–348 (2022).

36. Gao, G. P. et al. Novel adeno-associated viruses from rhesus monkeys as vectors for human gene therapy. Proc. Natl. Acad. Sci. U.S.A. 99 (18) 11854–11859 (2002).

37. Pillay, S. et al. An essential receptor for adeno-associated virus infection. Nature 530 (7588), 108–112 (2016).

38. Wang, D., Tai, P. W. L. & Gao, G. Adeno-associated virus vector as a platform for gene therapy delivery. Nat Rev Drug Discov. 18, pages 358–378 (2019).

39. Wang, J. H., Gessler, D. J., Zhan, W., Gallagher, T. L. & Gao, G. Adeno-associated virus as a delivery vector for gene therapy of human diseases. Signal Transduct. Target. Ther. 9,78, (2024).

40. Klein-Schneegans, A. S., Kuntz, L., Fonteneau, P. & Loor F. Serum Concentrations of IgM, IgGl, IgG2b, IgG3 and IgA in C57BL/6 Mice and their congenics at the lpr (lymphoproliferation) locus. J. Autoimmun. 2, 6, 869–875 (1989).

41. Swanson, S. J. What are clinically significant anti-drug antibodies and why is it important to identify them. Front. Immunol. 15, 1–8 (2024).

42. Wiswell, D. et al. A capillary electrophoresis based approach for the identification of anti-drug antibodies against camelid VHH biologics (Nanobodies ®). J Pharmacol Toxicol. Methods. 103 (2020).

43. Suarez-Amaran, L., Song, L., Tretiakova, A. P., Mikhail, S. A. & Samulski, R. J. AAV vector development, back to the future. Mol Ther. 33(**5**), 1903–1936 (2025).

44. Monteys, A. M. et al. Regulated control of gene therapies by drug-induced splicing. Nature 596, 7871, 291–295 (2021).

45. Desnoyers, L. R. et al. Tumor-Specific Activation of an EGFR-Targeting Probody Enhances Therapeutic Index. Sci. Transl. Med. 5, 207, (2013).

46. Labanieh, L. et al. Enhanced safety and efficacy of protease-regulated CAR-T cell receptors. Cell 185, 1745–1763 (2022).

47. Madeira, F. et al. The EMBL-EBI Job Dispatcher sequence analysis tools framework in 2024, Nucleic Acids Res 52, 1, 5, 521–525 (2024).

48. Beattie, J. et al. Molecular interactions in the insulin-like growth factor (IGF) axis: a surface plasmon resonance (SPR) based biosensor study. Mol Cell Biochem 307, 221–236 (2008).

49. Kalus, W. et al. Structure of the IGF-binding domain of the insulin-like growth factor-binding protein-5 (IGFBP-5): implications for IGF and IGF-I receptor interactions. EMBO J. 17, 22, 6558–6572 (1998).

50. Russo, V. et al. Granzyme B is elevated in autoimmune blistering diseases and cleaves key anchoring proteins of the dermal-epidermal junction. Sci Rep 8, 9690 (2018).

51. Hiroyasu, S. et al. Granzyme B inhibition reduces disease severity in autoimmune blistering diseases. Nat Commun 12, 302 (2021).

52. Darrah, E. & Rosen, A. Granzyme B cleavage of autoantigens in autoimmunity. Cell Death Differ. 17, 624–632 (2010).

53. Hull, C. M. et al. Granzyme B-activated IL18 potentiates αβ and γδ CAR T cell immunotherapy in a tumor-dependent manner. Mol. Ther., 32(**7**), 2373–2392 (2024).

54. Thompson, R., & Cao, X. Reassessing granzyme B: unveiling perforin-independent versatility in immune responses and therapeutic potentials. Front. Immunol. 15:1392535. (2024).

55. Rossana, T. et al. TGF-β Utilizes SMAD3 to Inhibit CD16-Mediated IFN-γ Production and Antibody-Dependent Cellular Cytotoxicity in Human NK Cells, J. Immunol. 181(**6**), 3784–3792 (2008).

56. Zhang, B., Zhang, J., & Tian, J. Comparison in the effects of IL-2, IL-12, IL-15 and IFN α on gene regulation of granzymes of human. Int. Immunopharmacol. 8, 989–996 (2008).

57. Mouchacca, P., Schmitt-Verhulst, A.M, & Boyer, C. Visualization of Cytolytic T Cell Differentiation and Granule Exocytosis with T Cells from Mice Expressing Active Fluorescent Granzyme B. PLoS ONE 8(**6**) (2013).

58. Choi, B.D., Yu, X., Castano, A.P. et al. CAR-T cells secreting BiTEs circumvent antigen escape without detectable toxicity. Nat Biotechnol 37, 1049–1058 (2019).

59. Fan, Y., Duan, Y., Chen, J., et al. Bispecific killer cell engager-secreting CAR-T cells redirect natural killer specificity to enhance antitumour responses. *Nat*. Biomed. Eng 10, 390–403 (2026).

60. Murad, J.P., Christian, L., Rosa, R. et al. Solid tumour CAR-T cells engineered with fusion proteins targeting PD-L1 for localized IL-12 delivery. *Nat*. Biomed. Eng 10, 613–629 (2026).

61. Radhakrishnan, H., Newmyer, S.L., Javitz, H.S., & P. Bhatnagar. Engineered CD4 T cells for in vivo delivery of therapeutic proteins. Proc. Natl. Acad. Sci. U.S.A. 121 (40) (2024).

62. Ping, Y., Li, F., Nan, S., Zhang, D., Shi, X., Shan, J., and Zhang, Y. Augmenting the Effectiveness of CAR-T Cells by Enhanced Self-Delivery of PD-1-Neutralizing scFv. Front. Cell Dev. Biol. 8(**803**) (2020).

63. S. Kan, S., et al. Delivery of an enzyme-IGFII fusion protein to the mouse brain is therapeutic for mucopolysaccharidosis type IIIB. Proc. Natl. Acad. Sci. U.S.A. 111 (41) 14870–14875, (2014).

64. Song, E.Z., Milone, M.C. Pharmacology of Chimeric Antigen Receptor–Modified T Cells. Annu. Rev. Pharmacol. Toxicol. 61, 805–829 (2021).

65. Freitas, K.A. et al. Enhanced T cell effector activity by targeting the Mediator kinase module. Science 378 eabn5647 (2022).

66. Guerrero, J.A., Klysz, D.D., Chen, Y., et al. GLUT1 overexpression in CAR-T cells induces metabolic reprogramming and enhances potency. Nat Commun 15, 8658 (2024).

67. Nimmerjahn, F. & Ravetch, J.V. Divergent Immunoglobulin G Subclass Activity Through Selective Fc Receptor Binding. Science 310, 1510–1512 (2005).

68. Sitaru, C., Mihai, S. & Zillikens, D. The relevance of the IgG subclass of autoantibodies for blister induction in autoimmune bullous skin diseases. Arch. Dermatol. Res. 299, 1–8 (2007).

69. Fridy, P. C. et al. A robust pipeline for rapid production of versatile nanobody repertoires. Nat. Methods 11, 1253–1260 (2014).

70. Han, J. et al. Relative Quantitation of Subclass-Specific Murine IgG Fc N-Glycoforms by Multiple Reaction Monitoring. ACS Omega 5 (**15**), 8564–8571(2020).

71. Kiyomi K., Muneo A., & Yuji M. A Simultaneous Semiquantification of Eight Immunoglobulin Isotypes in Mouse Serum by LC-MS/MS for Application of Immunogenicity Assessment of Therapeutic Antibodies. ACS Omega 10 (**22**), 23504–23513 (2025).

72. Maleitzke, T. et al. Protocol Standardized protocol and outcome measurements for the collagen antibody-induced arthritis mouse model Standardized protocol and outcome measurements for the collagen antibody-induced arthritis mouse model. STAR Protocols 3, 101718 (2022).

73. Lee, M.Y. et al. Collagen-Induced Arthritis Analysis in Rhbdf2 Knockout Mouse. Biomol. Ther. 26 (3), 298–305 (2017).

74. Crook, Z.R., Sevilla, G.P., Young, P., et al. CYpHER: catalytic extracellular targeted protein degradation with high potency and durable effect. Nat Commun 15, 8731 (2024).

75. Rhee, K., et al. GPCR Antagonism via Antibody-Mediated Rewiring of Receptor Trafficking and Degradation. Preprint at https://www.biorxiv.org/content/10.1101/2025.08.08.669350v1.full.pdf

76. Schett, G. et al. Advancements and challenges in CAR T cell therapy in autoimmune diseases. Nat. Rev. Rheumatol. 20, 531–544 (2024).

77. Kneitz, C., Wilhelm, M. & Tony, H. P. Effective B cell depletion with rituximab in the treatment of autoimmune diseases. Immunobiol. 206, 519–527 (2002).

78. Verdun, N. & Marks, P. Secondary Cancers after Chimeric Antigen Receptor T-Cell Therapy. *New Eng*. J. Med. 390 (**7**), 584–586 (2024).

79. Santiago-Ortiz, J. L. & Schaffer, D. V. Adeno-associated virus (AAV) vectors in cancer gene therapy. J. Control Release 240, 287–301 (2016).

80. Roybal, K. T. et al. Engineering T Cells with Customized Therapeutic Article Engineering T Cells with Customized Therapeutic Response Programs Using Synthetic Notch Receptors. Cell 167(**2**), 419–432 (2016).

81. Vlahos, A. E. et al. Protease-controlled secretion and display of intercellular signals. Nat. Commun. 13, 912 (2022).

82. Arjomandnejad, M., et al. Immunogenicity of Recombinant Adeno-Associated Virus (AAV) Vectors for Gene Transfer. BioDrugs 37, 311–329 (2023).

83. Byrne, B., et al. Current clinical applications of AAV-mediated gene therapy. Mol. Ther. 33, 2479–2516 (2025).

84. Muhuri, M. et al. Overcoming innate immune barriers that impede AAV gene therapy vectors. J Clin Invest. 131,1 (2021).

85. Srivastava, A., Mallela, K. M. G., Deorkar, N. & Brophy, G. Manufacturing challenges and rational formulation development for AAV viral vectors. J. Pharm. Sci. 110, 7, 2609–2624 (2021).

86. Shay, T. F. et al. Human cell surface-AAV interactomes identify LRP6 as blood-brain barrier transcytosis receptor and immune cytokine IL3 as AAV9 binder. Nat Commun. 15, 7853 (2024).

87. Hamilton, J. R. et al. In vivo human T cell engineering with enveloped delivery vehicles. Nat. Biotechnol. 42, 1684–1692 (2024).

88. Li, V. L. et al. An exercise-inducible metabolite that suppresses feeding and obesity. Nature 606, 785–790 (2022).

89. Yamada-Hunter, S.A., Theruvath, J., McIntosh, B.J., et al. Engineered CD47 protects T cells for enhanced antitumour immunity. Nature 630, 457–465 (2024).

90. Wickham, H. et al. Welcome to the Tidyverse, J. Open Source Softw. 4(**43**), 1686 (2019).

91. Barrett, T. et al. R package version 1.17.0; 10.32614/CRAN.package.data.table (2025).

92. Perez-Riverol, Y. et al. The PRIDE database at 20 years: 2025 update. Nucleic Acids Res. 53 (**D1**), D543–D553 (2025).

